# Molecular architecture of meiotic pro-crossover factor HEI10 reveals coupling of higher-order assembly and ubiquitin chain formation

**DOI:** 10.64898/2026.05.08.723602

**Authors:** Amy E. Milburn, Dhananjaya S. Kulkami, Carmen Espejo-Serrano, Miguel Pachon-Penalba, Moli E. Williams, Jonathan P. O. Nicol, Simona Debilio, Manickam Gurusaran, James M. Dunce, Ian R. Adams, Urszula L. McClurg, Neil Hunter, Owen R. Davies

## Abstract

In meiosis, crossovers between homologous chromosomes generate genetic diversity and are required for accurate chromosome segregation, ensuring fertility. In mammals, HEI10 is one of three pro-crossover RING-domain factors implicated in protein modification by ubiquitin and/or SUMO and characterised by their dynamic accumulation at future crossover sites. However, the molecular architecture and enzymatic activity of mammalian HEI10 have remained unknown. Here, we show that human HEI10 has E3-ubiquitin ligase activity that depends on its higher-order assembly. We report the crystal structure of the HEI10 core, revealing how a 29-nm rod-like tetramer is formed through head-to-head association of two coiled-coil dimers that results in clustering of four RING domains around the molecular centre. HEI10 tetramers self-assemble through RING, coiled-coil, and C-terminal interfaces into fibrous and spherical higher-order structures. Structure-guided mutants show that higher-order assembly is required for HEI10 to catalyse K63-linked ubiquitin chain formation *in vitro*, with the most active species likely corresponding to a loose, non-fibrous network of assembled HEI10 molecules. *Arabidopsis thaliana* HEI10 retains the tetrameric core and higher-order assembly behaviour, suggesting a conserved principle of HEI10 function.

## Introduction

Meiosis generates haploid gametes through a specialised programme of homologous chromosome pairing, synapsis, recombination and segregation (Hunter 2015; Zickler and Kleckner 2015). Central to the meiotic programme is the formation of crossovers between homologues through recombination. In addition to enhancing genetic diversity, crossovers establish physical connections between homologues that enable stable biorientation on the metaphase I spindle (Zickler and Kleckner 2023). Hence, crossovers are essential for meiosis, and reduced crossover formation in humans is associated with infertility, miscarriage and aneuploidy (Hassold and Hunt 2001; Nagaoka, Hassold, and Hunt 2012).

Meiotic recombination initiates in prophase I through the programmed formation of DNA double-strand breaks (DSBs) throughout the genome by SPO11-TOPOVIBL (Keeney, Giroux, and Kleckner 1997; Robert et al. 2016). Ensuing recombination-mediated repair utilises the homologue as the repair template (as opposed to the sister chromatid) to promote homologue pairing and end-to-end synapsis (Zickler and Kleckner 2023). In the context of synapsed homologues, a minority of recombination events – typically 10-30% - are matured into crossovers, with the remainder largely repaired as non-crossovers (Hunter 2015; Zickler and Kleckner 2015). This process is regulated to ensure that at least one crossover is formed between each homologue pair (crossover assurance), and that adjacent crossovers are widely and evenly spaced (crossover interference) (Gray and Cohen 2016; Wang et al. 2015). At a molecular level, most nascent strand-exchange intermediates (displacement-loops) are initially stabilised by MutSγ complex (MSH4–MSH5) (Snowden et al. 2004) and other meiosis-specific ZMM factors (Hunter 2015). This facilitates formation of the synaptonemal complex (SC) (Adams and Davies 2023; Llano and Pendas 2023; Moses 1956; Westergaard and von Wettstein 1972), a supramolecular protein structure that assembles between homologues and holds them together in a zipper-like manner along their entire length. MutSγ is subsequently lost from all sites other than prospective crossovers (Hunter 2015). Retention of MutSγ facilitates the formation of crossover-specific double-Holliday junctions (dHJs) and prevents their dissolution into non-crossovers until crossover-specific complexes, centred around the MutLγ (MLH1–MLH3) endonuclease, assemble and mediates resolution as crossovers (Kulkarni et al. 2020; Cannavo et al. 2020; Tang et al. 2025; Henggeler et al. 2025).

The crossover-specific retention of MutSγ requires CrossOver RING proteins (CORs) that share a similar domain organisation, with an N-terminal RING domain followed by a coiled-coil region and an unstructured C-terminus (Figure 1a,b). COR homologues have been identified widely across eukaryotes, but the number of paralogues varies from one in budding yeast, *Sordaria*, and *Arabidopsis* to four in *C. elegans* (Agarwal and Roeder 2000; Zhang et al. 2018; Nguyen et al. 2018; Bhalla et al. 2008; Chelysheva et al. 2012; Ziolkowski et al. 2017; Lake et al. 2015; Lake et al. 2019). Three CORs have been identified in mammals: RNF212, RNF212B and HEI10 (Ito et al. 2025). Cytologically, RNF212 and RNF212B behave similarly. In spermatocytes, they initially localise as numerous foci along nascent SCs before accumulating at crossover-designated sites while diminishing elsewhere (Reynolds et al. 2013; Condezo et al. 2024; Ito et al. 2025). In oocytes, RNF212-RNF212B still accumulate at crossover sites, but general loss from SCs is much less pronounced (Reynolds et al. 2013; Condezo et al. 2024; Ito et al. 2025). In spermatocytes, HEI10 forms prominent foci specifically at crossover-designated sites as RNF212-RNF212B accumulates (Qiao et al. 2014). In oocytes, prominent HEI10 foci are initially more numerous but eventually develop a crossover-specific pattern (Ito et al. 2025).

**Figure 1.**
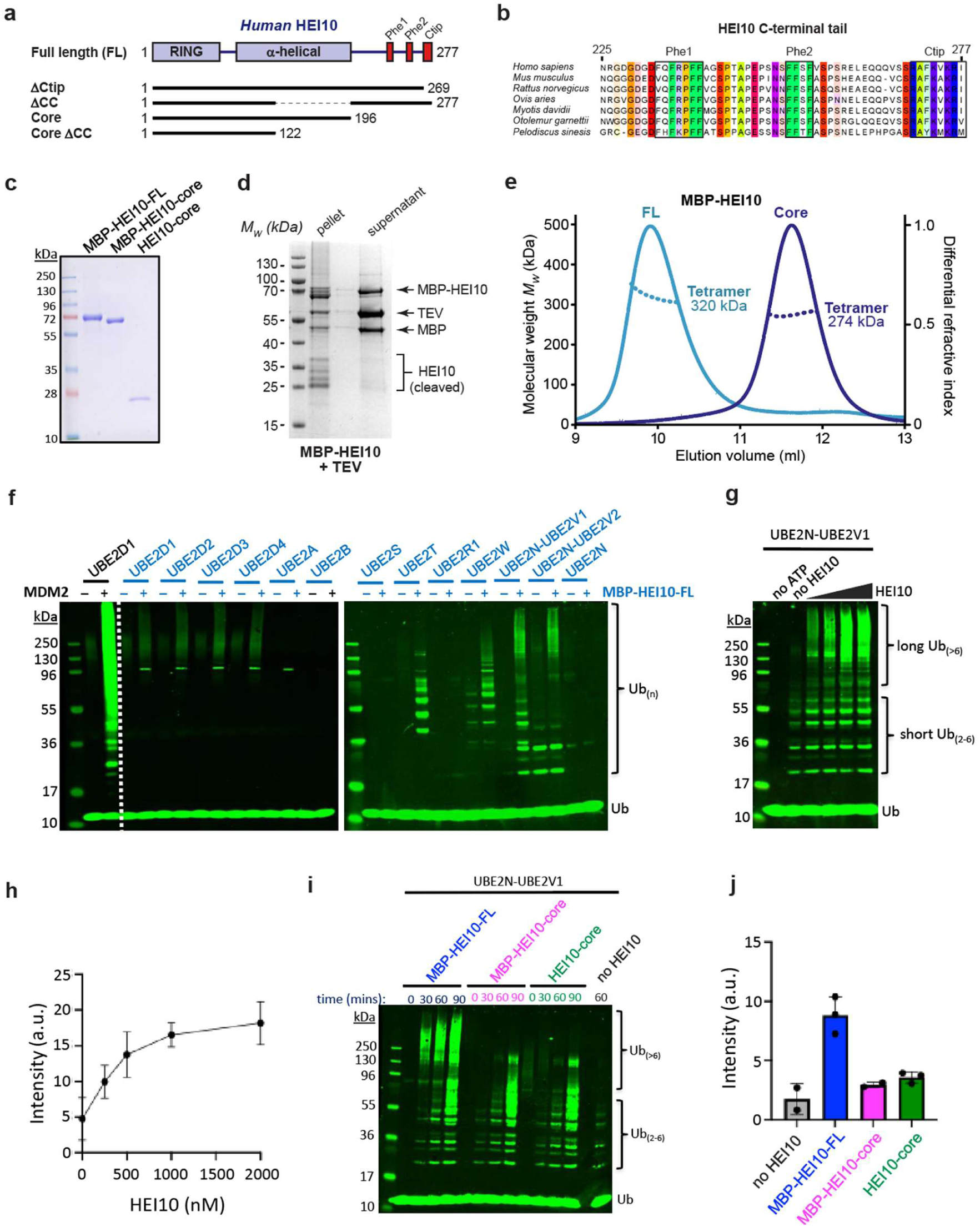
HEI10 is a tetramer that catalyses ubiquitin chain formation *in vitro*. (**a**) Schematic of the human HEI10 sequence depicting its RING domain, coiled-coil and the constructs used in this study. (**b**) Multiple sequence alignment of HEI10’s C-terminal tail, highlighting its conserved phenylalanine-rich patches (Phe1 and Phe2) and basic C-terminal tip (Ctip). (**c**) SDS-PAGE of purified MBP-HEI10-FL, MBP-HEI10-core and HEI10-core proteins. (**d**) The supernatant and pellet following TEV cleavage of the N-terminal MBP tag from HEI10-FL. (**e**) SEC-MALS analysis of MBP-HEI10, showing differential refractive index (dRI; solid lines) profiles with fitted molecular weights (*Mw*; dashed lines) across elution peaks. MBP-HEI10 FL (full-length; light blue) and core (amino-acids 1-196; dark blue) are 320 kDa and 274 kDa tetramers, respectively (theoretical – 311 kDa and 275 kDa). (**f**) Representative Western blot image of ubiquitylation assays for a panel of E2s with and without addition of MBP-HEI10-FL. MDM2 is shown as a positive control for E3-ligase activity. Ub, free ubiquitin; Ub_(n)_ ubiquitin chains. (**g**) Representative Western blot image of ubiquitylation assays with E2 UBE2N-UBE2V1 and varying concentrations of MBP-HEI10-FL (0.25 nM–2 μM). (**h**) Quantification of high molecular weight Ub_(>6)_ chains from the experiments represented in **g**. Means ± SEM for three independent experiments are shown. (**i**) Representative Western blot image of ubiquitylation time-courses with E2 UBE2N-UBE2V1 and either MBP-HEI10-FL, MBP-HEI10-core, or HEI10-core. (**j**) Quantification of high molecular weight Ub_(>6)_ chains from the experiments represented in **i** for the 60 minute time points. Means ± SEM for two or three independent experiments are shown.

Mutation of either RNF212 or RNF212B reduces the level of both proteins and strongly diminishes HEI10 focus formation. By contrast HEI10 mutation results in persistence of high numbers of RNF212 and RNF212B foci and stalling of recombination (Ito et al. 2025; Condezo et al. 2024; Qiao et al. 2014; Rao et al. 2017). Hence, it has been proposed that RNF212 and RNF212B stabilise MutSγ-associated recombination intermediates, while HEI10 promotes their selective turnover to differentiate crossover and non-crossover sites and allow recombination to progress (Qiao et al. 2014; Ito et al. 2025; Rao et al. 2017). Despite playing a central conserved role in crossing over, the molecular structure of the CORs and the mechanisms by which they regulate crossover formation remain unclear. CORs have long been implicated in SUMOylation and/or ubiquitylation (Rao et al. 2017; He et al. 2020; De Muyt et al. 2014; Cheng et al. 2006; Strong and Schimenti 2010; Wang et al. 2023), and evidence for E3-ligase activities has been presented for budding yeast Zip3 (He et al. 2021; Cheng et al. 2006), human HEI10 (Toby et al. 2003), and *Arabidopsis* HEI10 (Wang et al. 2023).

Here, we combine biochemistry, cell-based assembly assays and structural biology to define the molecular structure of human HEI10, the basis of its higher-order self-assembly, and the correlation with E3 ubiquitin ligase activity. HEI10 forms a stable tetrameric building-block that undergoes higher-order assembly through interfaces involving its RING domains, coiled-coil and C-terminal tail, which are important for HEI10 to stimulate the assembly of high-molecular-weight K63-linked ubiquitin chains. This core architecture is conserved in *Arabidopsis thaliana*, indicating evolutionary conservation of HEI10’s organisation. Together, our findings provide a structural framework for how pro-crossover RING proteins form higher-order functional assemblies to promote crossover formation during meiosis.

## Results

### HEI10 catalyses the formation of K63-linked ubiquitin chains in vitro

*In vitro* reconstitution of COR E3-ligase activities have been limited (Toby et al. 2003; Cheng et al. 2006; He et al. 2021) because the enzymes are largely insoluble and thus challenging to purify. HEI10 is comprised of an N-terminal RING domain followed by an α-helical coiled-coil and a C-terminal tail that is predicted to be unstructured but contains phenylalanine-rich patches and a basic C-terminal tip that are likely to mediate protein interactions (Figure 1a,b). While full-length human HEI10 (HEI10-FL) expressed in insect cells was insoluble, a HEI10-core construct (amino acids 1-196) was soluble indicating that insolubility is due to the unstructured C-terminal domain (Figure 1c,d). However, HEI10-FL was solubilised by fusion to an N-terminal MBP tag enabling purification to near homogeneity (Figure 1e). MBP-HEI10-FL formed a homogeneous tetrameric species as determined by size-exclusion chromatography multi-angle light scattering (SEC-MALS) (Figure 1e). MBP-HEI10-FL was screened for activity with a panel of E2 conjugases (UBE2s; Figure 1f). UBE2D-family (UBC5) E2s, which assemble K48-linked ubiquitin chains associated with proteasomal degradation, were only weakly stimulated by MBP-HEI10-FL. By contrast, MBP-HEI10-FL robustly stimulated several E2s that catalyse non-degradative ubiquitylation. UBE2T, also known as FANCT, is best known for its role in activating the Fanconi anaemia (FA) pathway of DNA-crosslink repair that interfaces with the recombination machinery in both somatic (Wilson et al. 2010; Raghunandan et al. 2015) and meiotic cells (Alavattam et al. 2016). UBE2T also influences Wnt/β-catenin, PI3K-AKT, and p53 signalling pathways (Gao et al. 2024). UBE2W uniquely catalyses α-amino N-terminal ubiquitylation (Tatham et al. 2013) that can limit target aggregation (Wang et al. 2018; Vittal et al. 2015), stabilise protein interactions (Scaglione et al. 2011), and provide an anchor for additional poly-ubiquitylation. For example, the tripartite motif (TRIM) family E3-ligase, TRIM21, employs this sequential mechanism to assemble self-anchored K63-chains that scaffold the activation of signalling kinases to drive the intracellular innate immune response (Foss et al. 2019; Fletcher et al. 2015). UBE2W has also been implicated in regulating the FA pathway (Zhang et al. 2011) UBE2N (UBC13) complexes, UBE2N-UBE2V1 and UBE2N-UBE2V2, pre-assemble short K63-linked ubiquitin chains that are subsequently transferred to substrates *en bloc* and help activate downstream pathways including the DNA-damage response, innate-immune and inflammatory signalling, and mitophagy (Hodge, Spyracopoulos, and Glover 2016; Lee et al. 2017; Pinder, Attwood, and Dellaire 2013; Geisler et al. 2014). For UBE2N-UBE2V1, stimulation of high molecular-weight K63-linked ubiquitin-chain formation by HEI10 was shown to be concentration dependent (Fig. 1g,h) and required all components of the ubiquitin machinery (E1, E2, E3, ubiquitin, and ATP; Supplementary Figure 1). Unexpectedly, the HEI10-core (with or without the MBP tag) lacked this activity (Fig. 1i,j), despite the fact that it still forms a homogeneous tetramer (Fig. 1e). Thus, sequences within the C-terminal tail are essential for HEI10 to function as an E3 ubiquitin ligase.

### HEI10 self-assembles into fibres and higher-order structures

Intriguingly, K63-linked ubiquitin chains can enhance protein accumulation and assembly (Hou and Tang 2023; Valentino et al. 2024; Dao et al. 2022; Galagedera et al. 2023), a process proposed to mediate assembly of amplified COR foci at designated crossover sites (Zhang et al. 2018; Morgan et al. 2021; Fozard, Morgan, and Howard 2023; Wang et al. 2023; Zhang et al. 2025). Further, the nature of HEI10 accumulation into prominent foci at crossover sites in mouse meiocytes suggests that higher-order self-assembly may be important for its function (Qiao et al. 2014; Ito et al. 2025). For these reasons, we examined the ability of human HEI10 to self-assemble into higher-order structures and their relationship with E3-ligase activity.

HEI10 became insoluble upon removal of the MBP tag by TEV cleavage (Figure 1d). Analysis of the insoluble material by negative-stain electron microscopy revealed that HEI10 self-assembled into fibres, in which individual fibrillar structures were typically 2-5 nm wide and up to 300 nm in length (Figure 2a and Supplementary Figure 2a). We also observed very large fibrous assemblies, of up to 150 nm in width and over 1 μm in length (Figure 2b and Supplementary Figure 2b). In all cases, fibres exhibited a striated appearance comprising longitudinal fibrils separated laterally by approximately 5 nm (Figure 2a,b and Supplementary Figure 2a,b). Hence, our observations are consistent with HEI10 assembling into a 2-5 nm fibrillar unit that undergoes lateral and longitudinal self-assembly to form higher-order fibrous structures.

**Figure 2.**
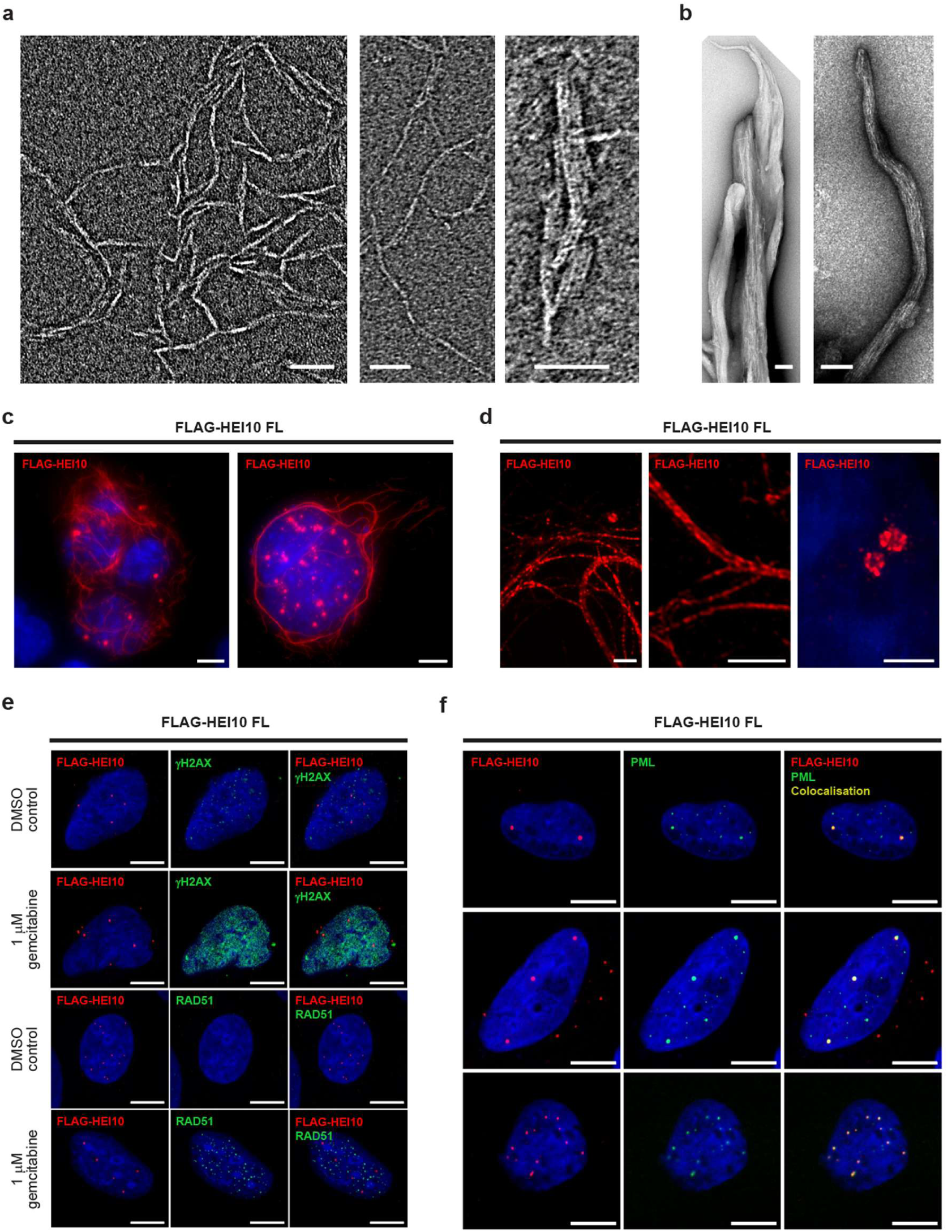
HEI10 self-assembly *in vitro* and *in vivo*. (**a,b**) Electron micrographs of HEI10 (full-length) pellets following TEV-cleavage from its N-terminal MBP tag, showing (**a**) internal fibrillar structure within HEI10 fibres, and (**b**) large fibrous assemblies. Scale bars, 50 nm in **a**; 100 nm in **b**. (**c,d**) Transient over-expression of Flag-HEI10 in COS7 cells, stained with anti-Flag antibody (red) and Hoechst 33342 (blue). (**c**) Wide-field imaging, showing that Flag-HEI10 forms cytoplasmic fibres and nuclear foci. Scale bars: 10 μm. (**d**) SoRa super-resolution imaging of cytoplasmic fibres (left) and nuclear foci (right). Scale bars: 2 μm. (**e,f**) Flag-HEI10 over-expression in U2OS cells. (**e**) Co-staining of Flag-HEI10 (red) with γH2AX (green; top) or RAD51 (green; bottom), and DAPI (blue), in cells treated with a DMSO control and 1 μM gemcitabine. Scale bars, 10 μm. (**f**) Co-staining of Flag-HEI10 (red), PML (green) and DAPI (blue), showing localisation of HEI10 foci in PML bodies. Scale bars, 10 μm.

As an orthogonal *in vivo* approach to analysing higher-order assembly, HEI10 was transiently expressed in COS7 cells. HEI10 formed both fibres and focal structures in this system (Figure 2c). HEI10 fibres were assembled almost exclusively in the cytoplasm, whilst HEI10 foci were seen in both the cytoplasm and nucleus (Figure 2c and Supplementary Figure 3a). Cytoplasmic fibres did not overtly colocalise with cytoskeletal markers, including tubulin, β-actin and cytokeratin (Supplementary Figure 4), suggesting that fibre formation may be an intrinsic property of HEI10 and not the result of association with pre-existing cellular structures. Further, SoRa super-resolution microscopy revealed an internal structure in which each cytoplasmic fibre consists of multiple closely associated parallel fibrils, consistent with their assembly through lateral and longitudinal interactions (Figure 2d and Supplementary Figure 3b). Hence, cytoplasmic HEI10 fibres formed *in vivo* may arise through the same intrinsic mechanism as the fibres observed *in vitro*.

The nuclear foci of HEI10 did not colocalise with sites of DNA damage marked by γH2AX or RAD51, either in untreated U2OS cells of in cells treated with gemcitabine, a DNA synthesis chain terminator (Figure 2e and Supplementary Figure 5a,b). Nuclear HEI10 foci were not associated with Cajal bodies or centrosomes (Supplementary Figures 6 and 7). However, almost all HEI10 nuclear foci co-localised with PML nuclear bodies (Figure 2f). We observed high Manders’ coefficient in both M1/HEI10 (0.8) and M2/PML (0.6). This indicates that most of HEI10 is localised to PML bodies, and a substantial proportion of PML bodies include HEI10. The remaining nuclear and cytoplasmic HEI10 foci may represent recruitment to other cellular structures or may simply represent smaller self-assembled structures. Together, these data indicate that HEI10 can self-assemble into fibres both *in vitro* and *in vivo* in the cytoplasm, and into foci that colocalise with PML nuclear bodies.

### HEI10 has a tetrameric core that self-assembles via its C-terminal tail

The solubility of the MBP-HEI10 core indicates that the C-terminal tail mediates the fibrous self-assembly that drives precipitation *in vitro*. To explore the structure of the tetrameric HEI10 core in solution (Figure 3a), SEC-MALS, circular dichroism (CD) and size-exclusion chromatography small-angle X-ray scattering (SEC-SAXS) were performed. SEC-MALS confirmed that the HEI10 core remains a homogeneous tetramer in the absence of MBP (Figure 3b). CD determined that the HEI10 core is approximately 50% α-helical, and is thermally stable, with a melting temperature of approximately 52°C (Supplementary Figure 8a,b). SEC-SAXS revealed a real-space *P(r)* interatomic distance distribution indicative of a rod-like molecule with a length of 29 nm (Figure 3c,d and Supplementary Figure 8c). Given that the predicted coiled-coil length within the core region is approximately 15 nm, the solution dimensions suggested that the tetramer comprises two coiled-coil segments arranged end-to-end. To define the minimal determinant for tetramer formation, we generated a construct that truncates the final 74 residues of the predicted coiled-coil (HEI10 core ΔCC; amino-acids 1–122; Figure 3a). SEC-MALS showed that HEI10 core ΔCC remained tetrameric, but with a markedly reduced molecular length of 11 nm determined by SEC-SAXS (Figure 3b-d and Supplementary Figure 8d), consistent with retaining the tetrameric unit whilst removing most of the rod-like extension. Further truncations beyond this point led to dissociation into smaller oligomers, indicating that the N-terminal portion of the coiled-coil is required for the integrity of the core tetramer. Finally, analysis using a spectrophotometric 4-(2-pyridylazo) resorcinol assay detected divalent cation binding for both HEI10 core and HEI10 core ΔCC proteins, consistent with zinc coordination by the RING domain (Supplementary Figure 8e). In summary, our biochemical data suggest that the building-block for HEI10 self-assembly is a core tetramer formed by its RING domains and the N-terminal end of the coiled-coils, which forms a 29-nm rod-like by virtue of the extended coiled-coils.

**Figure 3.**
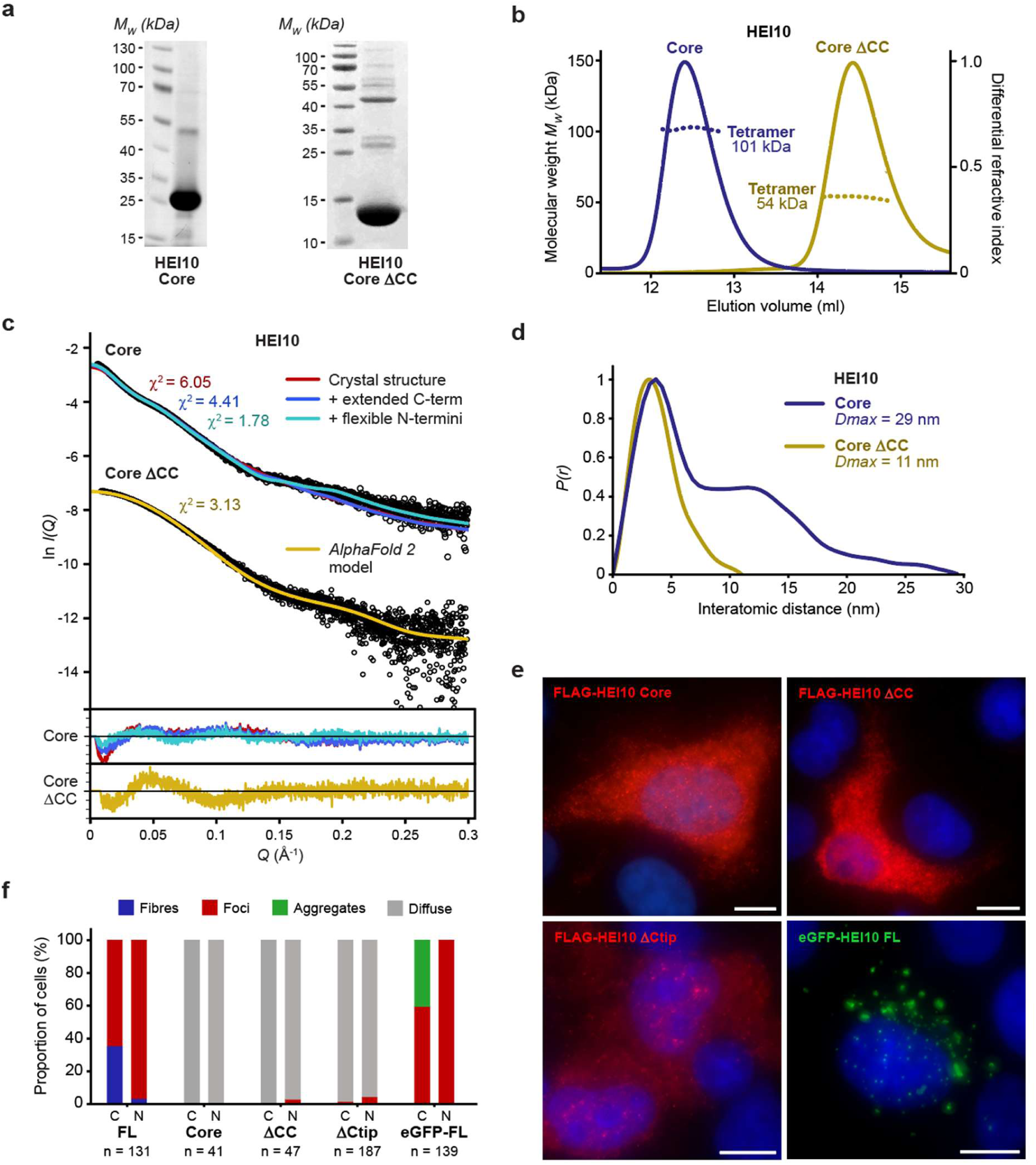
HEI10 has a tetrameric core that requires its C-terminal tail for self-assembly. (**a**) SDS-PAGE of purified HEI10 core (amino-acids 1-196) and core ΔCC (amino-acids 1-122). (**b**) SEC-MALS analysis showing that HEI10 core (blue) and core ΔCC (yellow) are 101 kDa and 54 kDa tetramers, respectively (theoretical – 97 kDa and 56 kDa). (**c,d**) SEC-SAXS analysis of HEI10 core and core ΔCC. (**c**) SAXS scattering data overlaid with the theoretical scattering curves of the HEI10 core crystal structure (red), with extended C-terminal coiled-coils (blue) and flexibly modelled N-termini (yellow), and with the HEI10 core ΔCC *AlphaFold 2* model (yellow); χ^2^ values are indicated and residuals for each fit are shown (inset). (**d**) SAXS *P(r)* interatomic distance distributions of HEI10 core (blue) and core ΔCC (yellow), showing maximum dimensions (*Dmax*) of 29 nm and 11 nm, respectively. (**e**) Transient over-expression of Flag-HEI10 ΔCtip (amino-acids 1-269), core (amino-acids 1-196) and core ΔCC (amino-acids 1-122) in COS7 cells, stained with anti-Flag antibody (red), and eGFP-HEI10, in COS7 cells. Scale bars, 10 μm. (**f**) Quantification of the presence of fibres (blue), foci (red), aggregates (green) and diffuse staining (grey) in the cytoplasm (C) and nuclei (N) of COS7 cells transiently transfected with HEI10 constructs, corresponding to imaging data shown in Figures 2a and 3e. The number of cells (n) is indicated for each construct.

We tested whether these structural features also govern higher-order assembly in cells. In agreement with its solubility *in vitro*, the HEI10 core failed to form fibres or foci upon transient expression in COS7 cells (Figure 3e,f). Notably, internal deletion of the coiled-coil segment corresponding to residues 123–196 (HEI10 ΔCC) also abrogated cellular fibre and foci formation (Figure 3e,f), despite this region not being required for tetramer formation *in vitro* (Figure 3b-d). Hence, the extended length of the coiled-coil — rather than tetramerisation *per se* — is necessary for higher-order assembly in cells. Further, an eight amino-acid truncation of the HEI10 C-terminal tip (HEI10 ΔCtip; amino-acids 1–269) blocked fibre formation, leaving only occasional nuclear foci, indicating that this conserved basic sequence is important for fibrous assembly (Figure 3e,f). Finally, fusion of HEI10 to an N-terminal eGFP tag blocked fibre assembly whilst retaining the ability to form cytoplasmic and nuclear foci (Figure 3e,f). Hence, the eGFP-HEI10 fusion functionally separates fibre and foci formation, indicating that these are related, but non-identical, processes. This behaviour is consistent with steric inhibition of fibril bundling by the N-terminal globular eGFP tag, providing a likely explanation for why recombinant full-length HEI10 is soluble as an N-terminal MBP fusion but precipitates upon tag removal (Figure 1d). Thus, our biochemical and cellular data indicate that HEI10 has a building-block rod-like tetrameric structure that assembles into fibres and foci via interactions mediated by its extended coiled-coil and C-terminal tail. Further, the formation of fibres, but not foci, may be sterically hindered by N-terminal fusion to a globular tag.

### Crystal structure of HEI10’s tetrameric core

To determine the molecular architecture of the HEI10 tetrameric building-block, we crystallised the HEI10 core. Crystals exhibited anisotropic diffraction to resolution limits between 2.4–5.7 Å (Table 1). The severe anisotropy hampered attempts at structure solution using the anomalous signal from bound zinc ions. Instead, we solved the structure by molecular replacement using an *AlphaFold 2* model of the HEI10 core DCC tetramer as a search model (Supplementary Figure 9). This provided sufficient phasing information to build the HEI10 core structure, in which amino-acids 8-184 were visible in electron density maps (Table 1 and Supplementary Figure 10a). The crystallographic model closely matches the *AlphaFold 2* model of the HEI10 ΔCC tetramer within the central region (r.m.s.d. = 0.94 Å; Supplementary Figure 10b), but the downstream coiled-coils diverge, explaining why structure solution was not achievable when using a model of the HEI10 core tetramer as a search model.

**Table 1.**
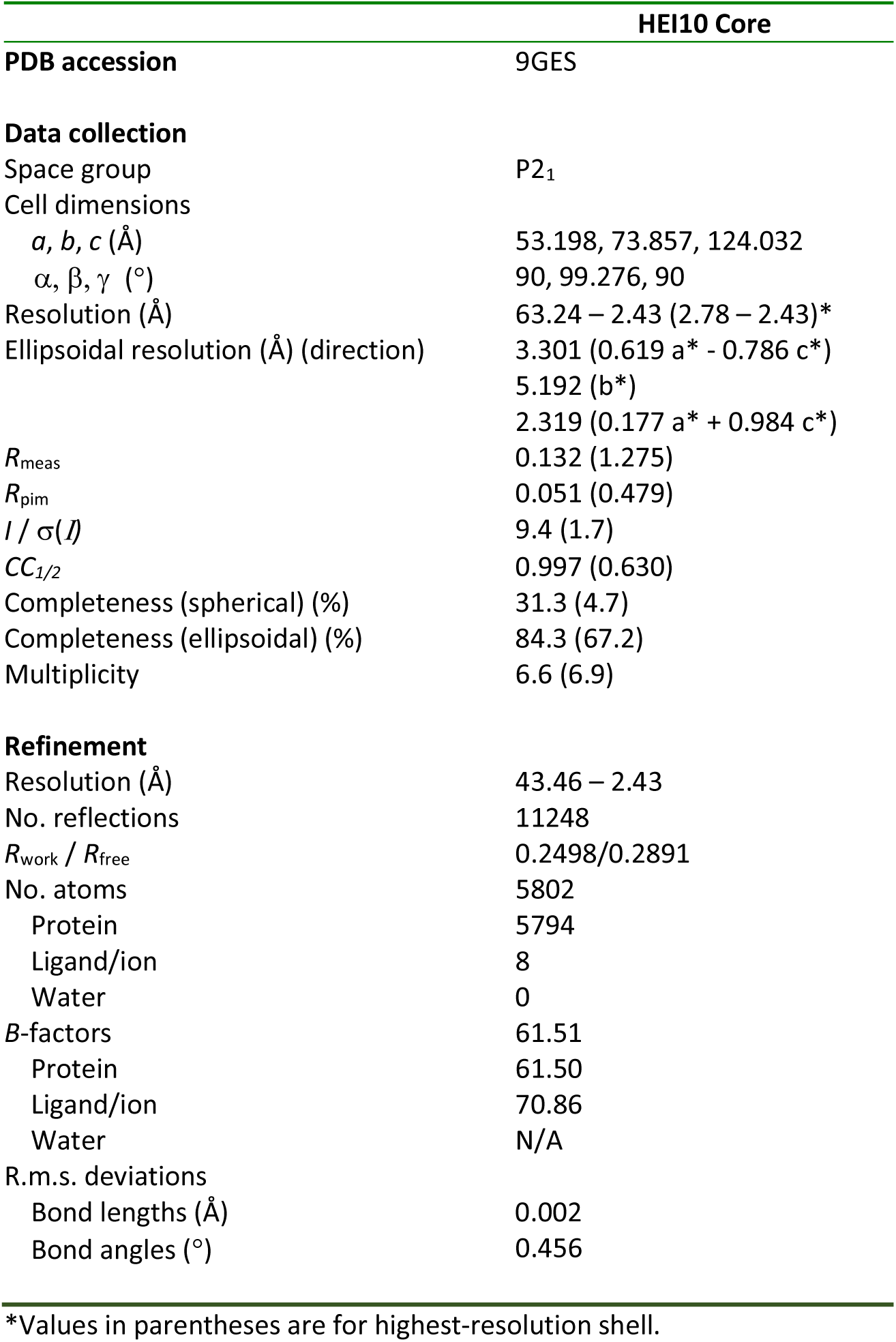
Data collection, phasing and refinement statistics.

|  | HEI10 Core |
| --- | --- |
| **PDB accession** | 9GES |
| **Data collection** |  |
| Space group | P2<sub>1</sub> |
| Cell dimensions |  |
| *a*, *b*, *c* (Å) | 53.198, 73.857, 124.032 |
| $\alpha$, $\beta$, $\gamma$ (°) | 90, 99.276, 90 |
| Resolution (Å) | 63.24 – 2.43 (2.78 – 2.43)* |
| Ellipsoidal resolution (Å) (direction) | 3.301 (0.619 *a* * - 0.786 *c* *) |
|  | 5.192 (*b* *) |
|  | 2.319 (0.177 *a* * + 0.984 *c* *) |
| *R*<sub>meas</sub> | 0.132 (1.275) |
| *R*<sub>pim</sub> | 0.051 (0.479) |
| *I* / $\sigma(I)$ | 9.4 (1.7) |
| *CC*<sub>1/2</sub> | 0.997 (0.630) |
| Completeness (spherical) (%) | 31.3 (4.7) |
| Completeness (ellipsoidal) (%) | 84.3 (67.2) |
| Multiplicity | 6.6 (6.9) |
| **Refinement** |  |
| Resolution (Å) | 43.46 – 2.43 |
| No. reflections | 11248 |
| *R*<sub>work</sub> / *R*<sub>free</sub> | 0.2498/0.2891 |
| No. atoms | 5802 |
| Protein | 5794 |
| Ligand/ion | 8 |
| Water | 0 |
| *B*-factors | 61.51 |
| Protein | 61.50 |
| Ligand/ion | 70.86 |
| Water | N/A |
| R.m.s. deviations |  |
| Bond lengths (Å) | 0.002 |
| Bond angles (°) | 0.456 |

\*Values in parentheses are for highest-resolution shell.

The HEI10 core crystal structure reveals a rod-like homo-tetramer of 29-nm length, in which the centre of the molecule is decorated by the four RING domains (Figure 4a). The tetramer is held together by a head-to-head interface between two coiled-coils (chains AC and BD), in which a four-helical bundle (amino-acids 82-96) has a hydrophobic core contributed by L86 and L93 side-chains from all four component monomers (Figure 4a). From this central interface, parallel dimeric coiled-coils extend C-terminally in opposite directions, giving rise to the overall 29-nm rod-like architecture (Figure 3a). At the N-terminal end of the coiled-coil, P82 caps the α-helical structure and packs against W96 from the opposing coiled-coil (Figure 4b). This configuration induces a sharp turn into a linker containing a short α-helix (amino-acids 70–75), which connects the coiled-coil to the RING domain (Figure 4c). The two linker helices from opposing chains interlock through interactions that include Y72, V76 and L77, and also bind to the underlying head-to-head interface (Figure 4a,c). This interlock enforces a 180° turn, positioning each RING domain against the downstream coiled-coil segment of its own chain (Figure 4a,c). Moreover, the RING domain is bound to its own coiled-coil through interactions that include amino-acids F95 and Y98 (Figure 4d,e). Hence, the four RING domains are tightly bound at the midline of the underlying coiled-coil structure to create the unusual molecular architecture of the tetrameric HEI10 core.

**Figure 4.**
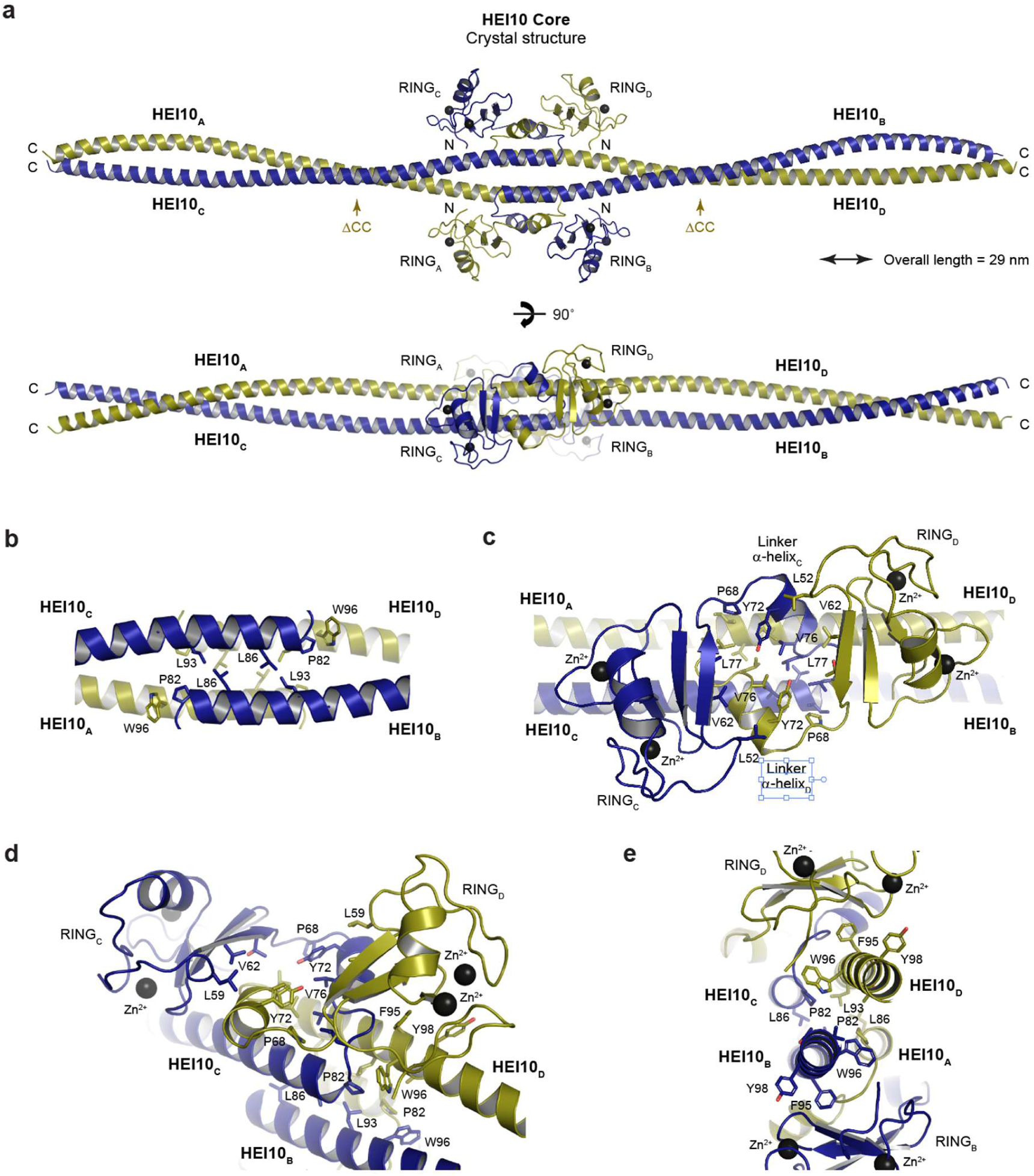
Crystal structure of the HEI10 core tetramer. (**a**) Crystal structure of HEI10 core (amino-acids 1-196). The rod-like homo-tetrameric assembly is 29 nm long and is formed through the head-to-head interaction of two parallel coiled-coils (in which constituent chains are coloured yellow and blue). At its centre, the N-terminal end of each coiled-coil chain is connected to the RING domain via a loop and short α-helix. The connecting α-helices of opposing chains interlock and chains turn back on themselves such that each RING domain binds to the downstream coiled-coil region of its own chain. The C-terminal boundaries of HEI10 ΔCC (amino-acids 1-122) are indicated by yellow arrows. (**b-e**) Structural details of the HEI10 core tetramer. (**b**) At the centre of the HEI10 structure, the head-to-head interface between opposing coiled-coils (chains AC and BD) is formed by a four-helical bundle with a hydrophobic core containing amino-acids L86 and L93 from all four chains. The α-helical coiled-coil structure is terminated at its N-terminal end by proline residue P82, which interacts with W96 of the opposing coiled-coil. (**c**) The N-terminal RING domain is connected to the coiled-coil via a short α-helix and loop. The short α-helices of opposing chains are interlocked in the midline through interactions that include amino-acids Y72, V76 and L77. (**d**) The interlocking short α-helices result in the RING domain of each chain packing against its own α-helical coiled-coil, stabilised by interactions including amino-acid F95. (**e**) Cross-section of the midline head-to-head interface, showing the roles of W96 and F95 in binding to the N-terminal end of the α-helical coiled-coil from the opposing chain and the RING domain from the same chain, respectively.

We confirmed that the HEI10 core crystal structure corresponds to the solution state by comparison with SEC-SAXS data. Firstly, the 29-nm length of the crystal structure agrees with the maximum dimension observed in the experimental SEC-SAXS real-space *P(r)* distribution (Figure 3d). Secondly, the SEC-SAXS scattering curve of the HEI10 core was closely fitted by the crystal structure upon modelling of short unstructured N-terminal sequences and C-terminal coiled-coil extensions that were present in the construct but not visible in electron density maps (χ² = 1.78; Figure 3c). We similarly analysed the ΔCC portion of the crystal structure that corresponds to the HEI10 core ΔCC construct (Figure 4a). This region predicts an 11-nm molecular length, matching the maximum dimension of the SEC-SAXS *P(r)* distribution, and its scattering curve was closely fitted by the corresponding structural model (*χ*^2^ = 3.13; Figure 3c,d). Thus, our crystallographic and solution data support a molecular architecture in which a head-to-head coiled-coil interface and interlocked linker α-helices constrain the RING domains into a compact central cluster around a rigid rod-like coiled-coil scaffold.

### Structural model for HEI10 self-assembly

How does the HEI10 tetrameric core assemble into the fibrous structures observed *in vitro* and in COS7 and U2OS cells? The HEI10 core crystal lattice is a fibrous array in which tetramers associate into linear fibrils that run in parallel and are separated laterally by 5 nm (Figure 5a). Within this lattice, linear fibrils are formed through longitudinal interactions along the length of staggered coiled-coils (Figure 5b), whereas lateral associations between fibrils are mediated by interactions between pairs of RING domains from tetramers in neighbouring fibrils (Figure 5c). Notably, this arrangement closely matches the striated appearance of full-length HEI10 fibres observed by electron microscopy, in which longitudinal fibrils are separated laterally by 5 nm (Figure 2a,b). Hence, we hypothesised that the HEI10 core crystal lattice defines the underlying architecture of HEI10 fibres, and that the full-length protein contributes additional interactions mediated by its C-terminal tail, resulting in stable fibres.

**Figure 5.**
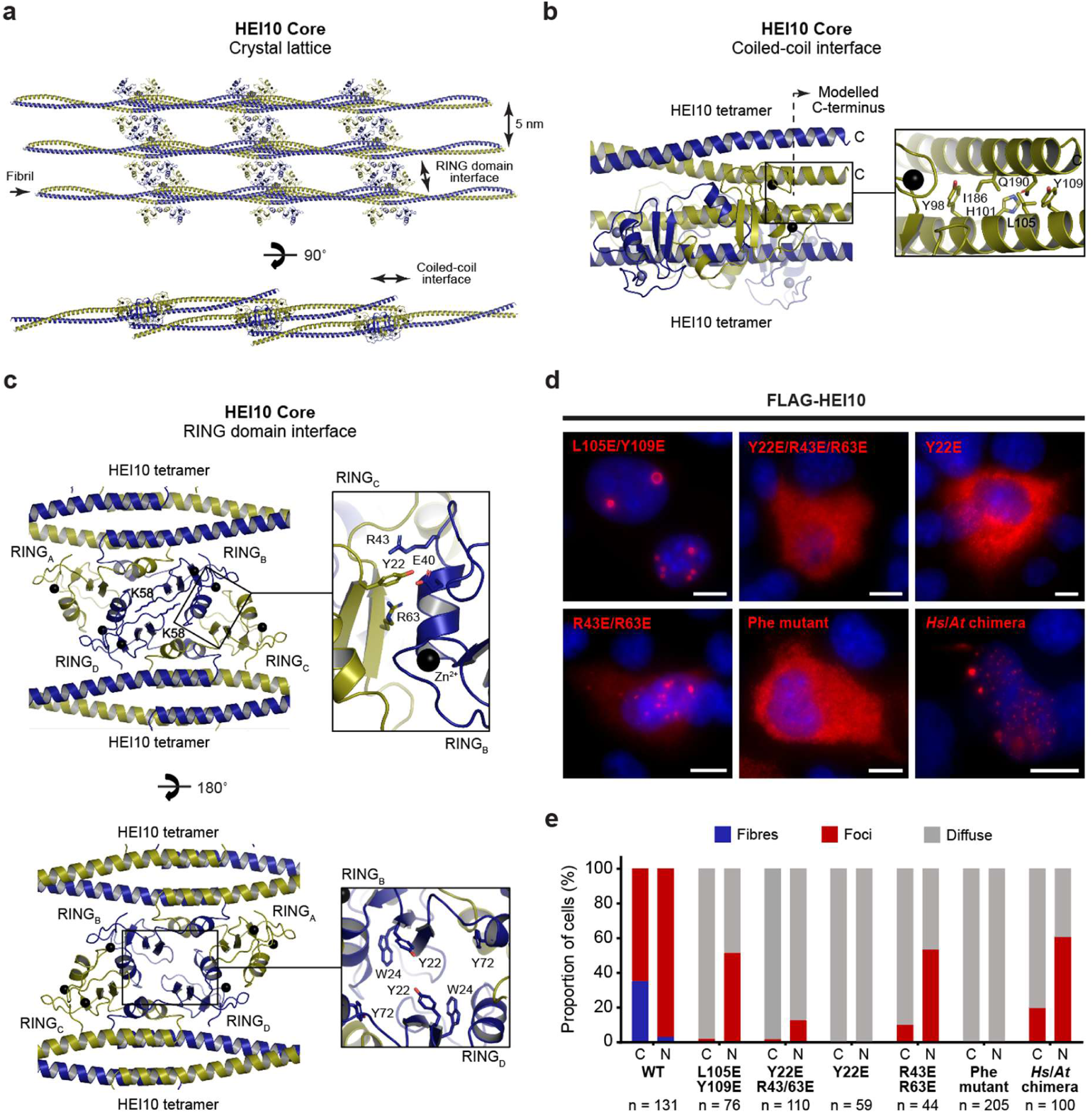
Molecular determinants of HEI10 self-assembly. (**a**) The HEI10 core crystal lattice consists of longitudinal fibrils separated laterally by 5 nm. Longitudinal fibrils are formed through anti-parallel interactions between HEI10 coiled-coils, whereas lateral packing is mediated by interactions between pairs of RING domains from parallel tetramers. The lower view is related to the upper view by a 90° rotation around the x-axis and a 15° rotation around the z-axis. (**b**) Longitudinal coiled-coil interface between the central head-to-head interface of one tetramer and the coiled-coil end of another. The α-helical coiled-coils have been extended beyond the amino-acids visible in the crystal structure (indicated by a dashed line), predicting that the interface is stabilised by interaction with amino-acids L105 and Y109. (**c**) The lateral interface between RING domains of parallel HEI10 tetramers is mediated by interactions between amino-acids Y22, R43 and R63 of opposing RING domains. The four RING domains form a tightly bound structure that is closed on one side (top), with a hollow cavity lined by aromatic amino-acids Y22, W24 and Y72 on its underside (bottom). (**d**) Wide-field imaging of over-expressed Flag-HEI10 constructs in COS7 cells, stained with anti-Flag antibody (red) and Hoechst 33342 (blue). Scale bars, 10 μm. The constructs analysed are mutants Y22E/R43E/R63E, Y22E, R43E/R63E and L105E/Y109E, glutamate mutations of the phenylalanine-rich patches (Phe mutant; F233E/F235E/F238E/F239E/F252E/F253E/F255E), and a chimeric protein (*Hs*/*At* chimera) in which the coiled-coil of human HEI10 (amino-acids 123-196) has been replaced by the equivalent coiled-coil from *Arabidopsis thaliana* HEI10 (amino-acids 113-184). (**e**) Quantification of the presence of fibres (blue), foci (red) and diffuse staining (grey) in the cytoplasm (C) and nuclei (N) of COS7 cells transiently transfected with HEI10 constructs, corresponding to imaging data shown in Figures 2a and 5e. The number of cells (n) is indicated for each construct.

To test this hypothesis, we designed separation-of-function mutants that target lattice interfaces whilst preserving the tetramer building-block structure. The longitudinal coiled-coil interface involves contacts between the C-terminal coiled-coil end of one tetramer and the central region of the coiled-coil from an adjacent tetramer (Figure 5a,b). In the context of an isolated tetramer, residues L105 and Y109 are solvent exposed, but in the crystal lattice they contribute to this longitudinal interface (Figure 5b). Accordingly, an L105E/Y109E mutation blocked fibre formation upon transient expression in COS7 cells, whilst retaining assembly of nuclear foci (Figure 5d,e). Thus, the longitudinal coiled-coil interface observed in the crystal lattice is required for HEI10 fibre assembly in cells.

The lateral packing between HEI10 fibrils is mediated by an inter-RING domain interface between tetramers in neighbouring fibrils, which includes contacts contributed by residues Y22, R43 and R63 (Figure 5c). This results in an intertwined RING domain tetramer, which is closed on its top surface by interlocked amino-acids K58 (Figure 5c, top panel), with a hollow interior lined by aromatic amino-acids Y22, W24 and Y72, and a wide opening on its underside (Figure 5c, bottom panel). Disruption of this interface by a Y22E/R43E/R63E triple mutation strongly abrogated HEI10 self-assembly, yielding only rare foci and no fibres (Figure 5d,e). The single Y22E mutation completely blocked both fibrous and focal assembly (Figure 5d,e), consistent with an essential role in self-assembly, either through its contribution to inter-RING binding or through shaping the aromatic-lined hollow cavity. In contrast, the R43E/R63E double mutant failed to assemble into fibres but retained nuclear foci (Figure 5d,e). Thus, similar to the N-terminal eGFP fusion and L105E/Y109E mutant, selective disruption of this lattice interface functionally separates fibre formation from foci assembly, indicating that ordered fibrous growth is sensitive to perturbation of the inter-RING domain packing interaction.

How might the HEI10 C-terminal tail contribute to these assemblies? The hollow cavity formed by the intertwined RING-domain tetramer is lined by aromatic residues (Figure 5c), creating a potential binding site for the conserved phenylalanine-rich patches within the C-terminal tail (Figure 1a). To test whether these motifs contribute to HEI10 assembly, we introduced glutamate substitutions across both phenylalanine-rich patches (Phe mutant; F233E/F235E/F238E/F239E/F252E/F253E/F255E). This mutant failed to form fibres or foci in cells (Figure 5d,e), indicating an essential role of the aromatic tail motifs in higher-order assembly. Together with the phenotype of HEI10 ΔCtip (Figure 3e,f), these data suggest that both the phenylalanine-rich patches and the basic C-terminal tip contribute to higher-order assembly of full-length HEI10.

In summary, our data support a structural model in which HEI10 fibres are built from a tetrameric rod-like core that polymerises through longitudinal coiled-coil interactions and lateral inter-RING domain packing interfaces that are captured in the HEI10 core crystal lattice, and are stabilised by conserved motifs within the C-terminal tail. Focal assembly is less sensitive to destabilisation of lattice interfaces, consistent with these representing looser networks of a less regular higher-order oligomer that is formed through the same self-assembly interfaces and also depends on the C-terminal tail (Figure 6a).

**Figure 6.**
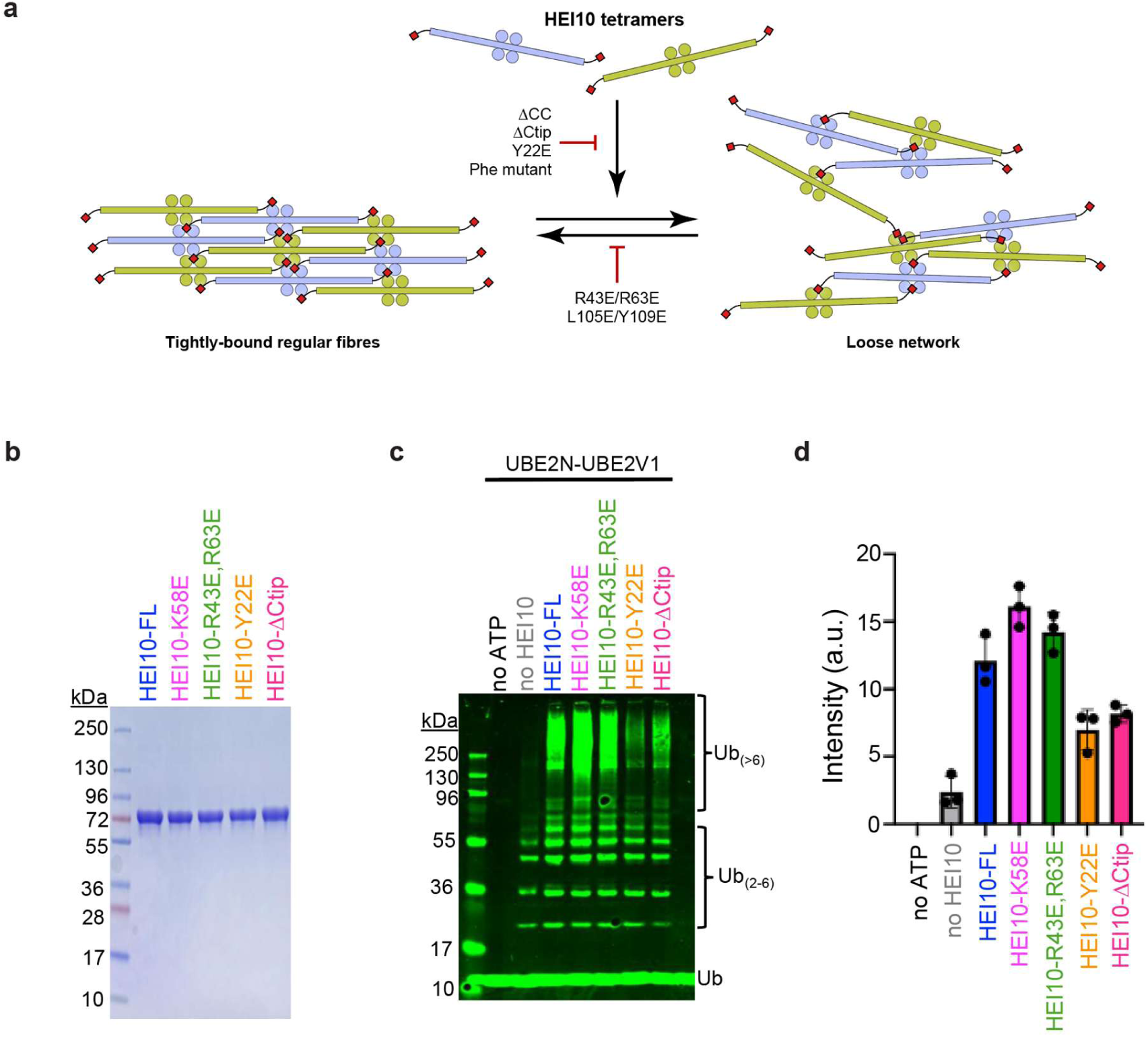
HEI10 ubiquitin ligase activity in mutants that alter higher-order assembly. (a) Schematic illustrating how HEI10 may self-assemble into tightly packed regular fibrous arrays and loose networks through common self-assembly interfaces. These consist of longitudinal coiled-coil contacts, inter-RING domain interactions and conserved motifs within C-terminal tails that stabilise higher-order structures. For clarity, only one C-terminal tail is illustrated for each dimeric coiled-coil. (b) SDS-PAGE of purified HEI10-FL, HEI10-K58E, HEI10-R43E,R63E, HEI10-Y22E and HEI10-ΔCtip proteins. All proteins are fused to a N-terminal MBP tag. (**c**) Representative Western blot image of ubiquitylation reactions with E2 UBE2N-UBE2V1 and HEI10 mutants shown in **b**. (**d**) Quantification of high molecular weight Ub_(>6)_ chains from the experiments represented in **c**. Means ± SEM for three independent experiments are shown.

**Figure 7.**
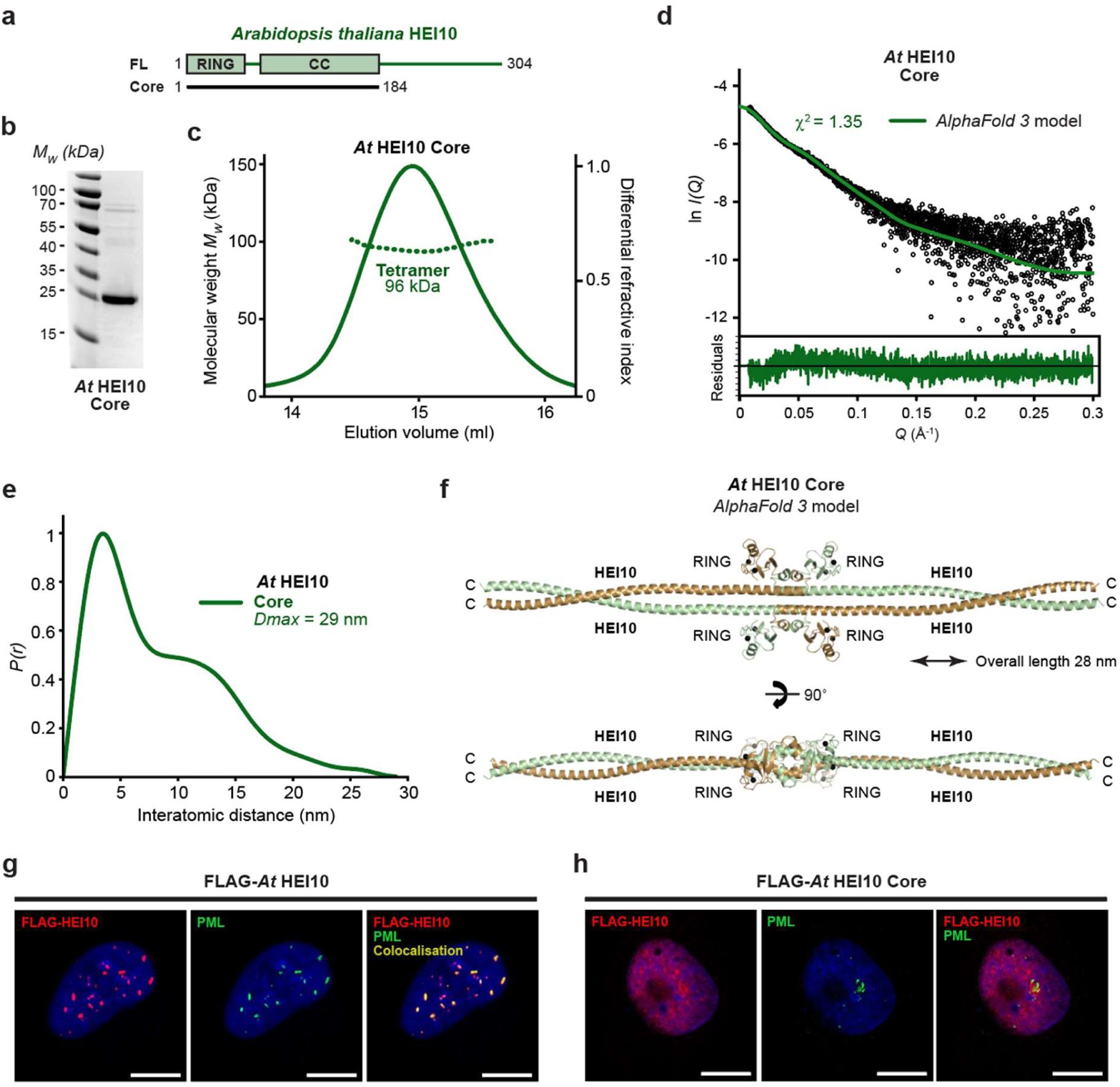
Structural model and self-assembly of *Arabidopsis thaliana* HEI10. (**a**) Schematic of the *Arabidopsis thaliana* HEI10 (*At* HEI10) sequence depicting its RING domain, coiled-coil and the constructs used in this study. (**b**) SDS-PAGE of purified *At* HEI10 core (amino-acids 1-184). (c) SEC-MALS analysis showing that *At* HEI10 core is a 96 kDa tetramer (theoretical – 92 kDa). (**d**,**e**) SEC-SAXS analysis of *At* HEI10 core. (**d**) SAXS scattering data overlaid with the theoretical scattering curve of the *AlphaFold 3* model (green); χ^2^ values are indicated and residuals for each fit are shown (inset). (**e**) SAXS *P(r)* interatomic distance distribution, showing a maximum dimension (*Dmax*) of 29 nm. (**f**) *AlphaFold 3* model of the *At* HEI10 core tetramer, showing the same architecture as the human HEI10 core structure, with an overall length of 28 nm. (**g**,**h**) Wide-field imaging of over-expressed Flag-*At* HEI10 (**g**) full-length and (**h**) core constructs in U2OS cells, with co-staining of Flag-HEI10 (red), PML (green) and DAPI (blue), showing localisation of full-length but not core HEI10 foci to PML bodies. Scale bars, 10 μm.

### HEI10 higher-order self-assembly correlates with E3 ubiquitin ligase activity

Our observation that the HEI10 core lacks E3-ligase activity (Figure 1i), led us to further explore the correlation between higher-order assembly and enzymatic activity (Figure 6b-d). HEI10 mutant variants were purified to near homogeneity (Figure 6b; all variants are N-terminal MBP fusions) and tested for *in vitro* ubiquitylation activity with UBE2N-UBE2V1 (Figure 6c,d). K58E was hyperactive for E3-lligase activity (Figure 6c,d) suggesting that disrupting the interlocked surface of the inter-RING interface can enhance activity. The R43E/R63E double mutant was also hyperactive, though less so than K58E (Figure 6c,d). Thus, E3-ligase activity is positively correlated with the ability of HEI10 to assemble foci but not fibers. This inference was supported by analysis of the Y22E and ΔCtip mutants, both of which abolish both fibre and focus assembly and had diminished E3-ligase activity (Fig. 6c,d). Collectively, these data support a model in which loosely assembled HEI10 foci activate E3-ligase activity, while highly-ordered fibres are inhibitory.

### Conservation and divergence of the functional architecture of HEI10 in Arabidopsis thaliana

Localisation of mammalian HEI10 to crossover sites is dependent on two other pro-crossover RING-proteins, RNF212 and RNF212B (Reynolds et al. 2013; Qiao et al. 2014; Rao et al. 2017; Ito et al. 2025). By contrast, *Arabidopsis thaliana* (*At*) encodes a single pro-crossover RING protein, HEI10, and lacks orthologues of RNF212 and RNF212B. Its sequence is conserved relative to mammalian HEI10 across the structured N-terminal core (RING domain and coiled-coil), but diverges substantially within the C-terminal tail, lacking the mammalian phenylalanine-rich motifs and instead containing distinct motifs that are conserved within plants (Figure 7a). We therefore asked whether *At* HEI10 adopts the same tetrameric building-block architecture as human HEI10, and whether it exhibits a similar propensity for higher-order assembly into fibres and/or foci.

To define *At* HEI10’s core architecture, we purified an *At* HEI10 core construct (amino-acids 1–184; Figure 7b). The recombinant protein was soluble, and SEC-MALS revealed a single dominant peak corresponding to a monodisperse tetramer (Figure 7c). SEC-SAXS analysis indicated an elongated structure with a real-space *P(r)* distribution consistent with a rod-like molecule of 29 nm length (Figure 7d,e and Supplementary Figure 11). Thus, *At* HEI10 core has the same oligomeric state and dimensions as the human HEI10 core tetramer. An *AlphaFold 3* model of the *At* HEI10 core tetramer predicts the same overall architecture as human HEI10, in which a head-to-head association of parallel dimeric coiled-coils clusters the four RING domains at the midline (Figure 7f and Supplementary Figure 12). The model further predicts conservation of the linker-mediated packing of RING domains against the coiled-coil, including interlocked linker helices (Figure 7f). Although the predicted coiled-coil arrangement likely contains local inaccuracies, the *AlphaFold 3* model closely fitted the experimental SEC-SAXS scattering data (χ^2^ = 1.35; Figure 7d), supporting the conclusion that its global architecture is correct. Hence, the tetrameric core building-block of HEI10 appears to be conserved between mammals and plants.

Upon transient expression in U2OS cells, *At* HEI10 formed prominent foci that were predominantly nuclear, but we did not observe the cytoplasmic fibres that were readily formed by human HEI10 (Figure 7g). Thus, focal assembly is conserved in this heterologous system, whereas fibrous assembly is not. This agrees with previous reports (Wang et al. 2023). Consistently, a chimeric human HEI10 in which the human coiled-coil was replaced by the corresponding *At* HEI10 coiled-coil retained the ability to form foci but failed to assemble into fibres (Figure 5d,e), indicating that fibrous assembly depends on sequence features that are specific to mammalian HEI10 rather than being an inevitable consequence of the conserved tetrameric architecture. Co-staining showed that a substantial fraction of large *At* HEI10 foci co-localised with PML (Figure 7g and Supplementary Figures 13 and 14), indicating that recruitment to PML bodies is conserved despite evolutionary divergence. Finally, deletion of the C-terminal tail (*At* HEI10 core; amino-acids 1–184) largely disrupted focus formation, restricting HEI10 to a diffuse nuclear staining pattern (Figure 7h and Supplementary Figure 15). Thus, as in mammals, *At* HEI10 focal assembly requires the C-terminal tail, despite the absence of phenylalanine-rich motifs that are important in human HEI10, suggesting functional conservation through sequence divergence.

In summary, *At* HEI10 adopts a conserved tetrameric, rod-like building-block architecture and undergoes C-terminal tail-dependent focal assembly in cells. However, *At* HEI10 does not form fibres, and substitution with the plant coiled-coil blocks fibrous assembly by human HEI10. Hence, whilst the HEI10 tetramer is evolutionarily conserved, the modes of higher-order assembly and the molecular interactions that drive them have diverged, potentially reflecting differences in how plant and mammalian HEI10 are localised to crossover sites during meiosis.

## Discussion

The molecular structures of meiotic pro-crossover RING proteins and their relationships with enzymatic activities had remained unknown. Here, we provide evidence that human HEI10 is an assembly-dependent ubiquitin E3-ligase built around a tetrameric core. The HEI10 tetramer is a unique 29-nm rod-like structure in which two parallel coiled-coil dimers associate head-to-head, clustering the four RING domains at the centre and extending C-terminally in opposite directions. HEI10 self-assembles into fibres and foci through a common mechanism, involving longitudinal coiled-coil contacts, lateral inter-RING interactions and conserved motifs within its C-terminal tail that stabilise higher-order structures. The fibres formed *in vitro* and in cells likely represent an unrestricted state in which these interfaces are fully engaged to generate tightly packed, regular molecular arrays. By contrast, cellular foci likely represent looser, irregular and more solvent-accessible networks formed through the same interactions, but with partial interface occupancy. Consistent with this model, mutants that weaken assembly interfaces disrupt fibres while preserving foci, whereas mutants that abolish these interfaces prevent both fibres and foci, restricting HEI10 to its unassembled tetrameric state. Thus, HEI10 tetramers use a common mechanism to generate higher-order assemblies ranging from regular fibres to loose networks (Figure 6a).

We find that HEI10 stimulates ubiquitylation in reconstituted reactions by several E2s. Focusing on the assembly of high molecular-weight K63-linked ubiquitin chains by UBE2N-UBE2V1/2, we showed that stimulation of this activity by HEI10 requires its disordered C-terminal tail and correlates with the ability to assemble foci but not fibres *in vivo*. Indeed, mutants that block fibre formation but retain looser focal assemblies are hyperactive, indicating that the most ordered state is not the most catalytically productive. Hence, regular fibres may be an unrestricted endpoint that forms readily *in vitro* but limits catalysis owing to steric hindrance, whereas loose networks may enable interaction with E2 enzymes, so are more active. This model is consistent with the extended length of the HEI10 coiled-coil, which is dispensable for tetramer formation but required for assembly of fibres and foci.

In addition to driving assembly, the coiled-coil may space RING domains within higher-order structures to preserve E2 access and coordinate activity between HEI10 subunits. Whilst the physiological substrates of human HEI10 remain unclear, the ability to catalyse non-degradative K63-linked chains suggests a role in assembly and/or signalling as opposed to proteolysis. These activities of human HEI10 are consonant with the inference that *Arabidopsis* HEI10 self-associates and promotes self-association of other crossover factors (Wang et al. 2023). However, *Arabidopsis* HEI10 is also implicated in ubiquitylation and degradation of AtRPA1a, an *Arabidopsis* RPA1 paralog that is important for crossing over (Wang et al. 2023). We observed only weak stimulation of K48-linked chains by UBE2D-family E2s (Fig. 1f). However, HEI10 might stimulate other K48-chains by other E2s or under different conditions. Alternatively, HEI10-catalyzed K63 chains may indirectly lead to degradation by seeding the formation of branched K63/K48 chains (Ohtake et al. 2018)

Upon expression in somatic cells, HEI10 was recruited to PML nuclear bodies in a manner that depended on residues required for higher-order assembly. PML nuclear bodies (NBs), membrane-less organelles implicated in regulating a variety of cellular processes (Lallemand-Breitenbach and de The 2018; Rerolle and de The 2023; Abou-Ghali and Lallemand-Breitenbach 2024; Corpet et al. 2020; Lallemand-Breitenbach and de The 2010). Nascent PML NBs organize polyvalent interaction shells, comprising both the SUMO-interaction motif (SIM) of PML and SUMO conjugated to PML, that recruit client proteins modified by SUMO and/or containing SIMs. These include the SUMO machinery, which further modifies clients to reinforce PML NB localisation and regulate their stability and activities. HEI10 also binds SUMO (Strong and Schimenti 2010) and can be sumoylated *in vitro* (unpublished). HEI10 may be SUMO modified *in vivo* and/or may contain active SIMs, as suggested by a previous report that HEI10 binds SUMO in yeast two-hybrid assays (Strong and Schimenti 2010). Thus, we suggest that HEI10 accumulation at both PML NBs in the orthogonal cell system, and at designated crossover sites during meiosis, may be driven by higher-order assembly and SUMO-SIM interactions. Suggestively, the other mammalian CORs, RNF212 and RNF212B, are inferred to catalyse sumolyation and required for efficient localization to crossover sites (Rao et al. 2017; Ito et al. 2025).

The finding that HEI10 forms tetramers and distinct higher-order assemblies raises the question of which of these states operate in meiosis. We suggest that HEI10 assembly may be regulated spatially and temporally during prophase, such that different structures are favoured at different stages and chromosomal sites. Transitions between these states could be driven by protein interactions and/or post-translational modification. Analysis of HEI10 association with the SC in mouse spermatocytes (Qiao et al. 2014) suggests that mutually reinforcing effects of recombination, synapsis and developing crossover sites could modulate its assembly state. Cytologically, HEI10 is most prominent as discrete foci at designated crossovers (Qiao et al. 2014; Ito et al. 2025), a pattern more consistent with local loose networks than with extended fibres. However, genetic evidence indicates that HEI10 acts before these bright foci become visible, implying an earlier, cytologically undetectable pool (Qiao et al. 2014). Possibly, HEI10 initially assembles into catalytically inactive fibres or fibrillar arrays early in meiosis, either along the SC or in non-chromosomal pools. As meiosis progresses, these structures could be remodelled and concentrated into loose networks of catalytically active HEI10 at designated crossover sites. A plausible regulator of HEI10 localization and assembly transitions is RNF212/RNF212B-dependent sumoylation. These proteins may either modify HEI10 directly or generate a SUMO-rich environment that promotes its recruitment, accumulation and catalytic activity at designated crossover sites.

The conservation of the tetrameric scaffold in *Arabidopsis thaliana*, together with its ability to form foci in cells, suggests that regulated higher-order assembly is a conserved feature of HEI10 function. Defining the endogenous ultrastructural state of HEI10, and how SUMO, RNF212 and RNF212B shape its assembly and activity, will be crucial for understanding how supramolecular organisation controls crossover formation during meiosis.

## Materials and Methods

### Purification of human HEI10 for ubiquitylation assays

Human HEI10 coding region was synthesized (GeneCopoeia) and cloned into the modified pFastBac (6His-MBP-TEV pFastBac vector, a kind gift from Jawdat Al-Bassam, UC Davis). The resulting 6His-MBP-TEV-HEI10 construct was expressed in Sf-9 cells. Baculovirus was infected at 1x10^6^ per mL and cells were incubated at 27°C for 52 hrs with shaking. Cells were centrifuged at 1500 rpm for 10 minutes at 4°C, washed with ice cold PBS, frozen in liquid nitrogen and stored at −80°C. A typical purification was performed with cell pellets from 800 ml of culture. Cells were resuspended in 4 volumes of lysis buffer (50 mM HEPES-NaOH pH 7.4, 1 mM DTT, 1 mM EDTA, 1:500 (v/v) protease inhibitor cocktail (P8340 Sigma), 1 mM phenylmethylsulfonyl fluoride, 20 μg/ml leupeptin) and stirred slowly for 15 min. Glycerol was added to 16% and 5 M NaCl was added to a final concentration of 325 mM. The sample was stirred for additional 20 min and then centrifuged at 30,000 g for 30 min. The clarified extract was bound in batch mode with 3 ml of pre-equilibrated amylose resin (New England Biolabs) for 1 hr. The resin was then washed extensively with 200 ml of buffer A (50 mM HEPES-NaOH pH 7.4, 1 mM β-DTT, 300 mM NaCl, 10 % glycerol, 1 mM phenylmethylsulfonyl fluoride, 10 μg/ml leupeptin). Bound protein was eluted with the above buffer containing 10 mM maltose. Peak fractions were collected and further purified using a Superdex200 10/300GL (Cytiva) gel-filtration chromatography column and buffer containing HEPES-NaOH pH 7.4, 1 mM β-DTT, 300 mM NaCl, 10 % glycerol, 1 mM phenylmethylsulfonyl fluoride. The peak fractions were pooled and dialyzed with buffer B containing HEPES-NaOH pH 7.4, 1 mM β-DTT, 100 mM NaCl, 10 % glycerol, 1 mM phenylmethylsulfonyl fluoride, and then flash frozen in liquid nitrogen and storage at −80 °C.

Human HEI10 mutants (hHEI10 Y22E, K58E, R43E/R63E double mutant, and hHEI10 1-269) were costructed using a QuikChange II Site-Directed Mutagenesis Kit (Agilent Technologies Inc, 200524). All HEI10 mutant proteins were expressed as His-MBP fusions and prepared as described above. The His-MBP tag was cleaved from His-MBP-HEI10-core by Hi-TEV protease and both the cleaved tag and protease (His-MBP and His-TEV) where removed by passing through a Ni-NTA column. Tag-less HEI10-core was concentrated and further purified by gel-filtration chromatography column as described above. All the protein concentrations were determined by spectroscopic absorption at 280 nm using extension coefficients and molecular weights.

### Purification of Ubiquitin

Ubiquitin was expressed in *E. coli* (BL21 DE3 cells). A plasmid containing GST-TEV-FLAG-Ubiquitin (a kind gift from Nitzen Shabek lab UC Davis), was grown in 2xYT media at 37 °C to an OD_600_ of 0.8, then expression was induced with 1 mM ITPG for 16 hr at 20 °C. Induced cells were harvested by centrifugation, washed with PBS buffer and lysed by sonication (50 amplitude, 20s on/30s off cycle for 5 min) on an ice bath in buffer A (50 mM Tris pH 8.0, 500 mM NaCl, 10% Glycerol, 0.5 mM EDTA, 1 mM PMSF, protease inhibitors cocktail - Roche, 04693159001). The lysate was centrifuged at 30,000 rpm for 30 minutes at 4 °C using a Ti45 rotor (Beckman Coulter). The supernatant was applied to a GST affinity column, pre-equilibrated with buffer A, which was then washed with low salt buffer B (50 mM Tris pH 8.0, 25 mM NaCl, 5 % Glycerol, 1 mM EDTA, 1 mM PMSF). The GST tag was cleaved with TEV protease on the column overnight. The flow was collected and injected onto a 1 mL HiTrap Q HP (Cytiva) ion exchange chromatography column equilibrated with buffer B. The column was washed with buffer B and bound proteins were eluted with a salt gradient. The peak fractions collected were pooled and concentrated, flash frozen in liquid nitroge,n and storage at −80 °C. Protein concertation was determined as described above.

### Ubiquitylations Assays

Ubiquitin conjugation reactions were performed under multi-turnover conditions at 37 °C in 15 uL reactions containing 20 mM HEPES, pH 7.5, 50 mM NaCl, 5 mM ATP, 5 mM MgCl_2_, 0.01% Tween and 0.5 mM DTT; and 25-75 nM E1 (UBA1, Boston Biochem), 25-100 nM E2 conjugating enzymes (UBE2s, from UBPbio), 10 uM FLAG-Ubiquitin and indicated concentrations of HEI10. Reactions were terminated by addition of Nu-PAGE LDS sample loading buffer (Invitrogen, NP0007) with reducing agent, and separated on Nu-PAGE 4-12% Bis-Tris Gels (Invitrogen NP0336B) at 120V with MES-SDS running buffer (Invitrogen, NP0002), followed by Western blotting using anti-FLAG antibodies (Sigma-Aldrich F1804) to detect FLAG-ubiquitin. Blots were imaged using an Odyssey Infrared Imager (LI-COR) and quantified using image J software version 1.53k. All the other recombinant enzymes used in this study were obtained from Boston Bioche or Sigma-Aldrich.

### Recombinant protein expression and purification for structural analysis

Sequences corresponding to regions of human HEI10 and *Arabidopsis thaliana* HEI10 (*At* HEI10), were cloned into pRSF-Duet1 (Novagen®) expression vectors for expression as TEV-cleavable N-terminal His_6_- and MBP-fusion proteins, as described. Constructs were co-expressed in Rosetta2 (DE3) cells (Novagen®), in Terrific Broth media supplemented with 50 µM ZnOAc, induced with 0.5 mM IPTG for 16 hours at 25°C. Cells were lysed by sonication in 50 mM Tris pH 8.0, 500 mM NaCl, 10% glycerol, 50 µM ZnOAc, 5 mM DTT, and fusion proteins were purified from clarified lysate through consecutive amylose (NEB) affinity chromatography and HiTrap Q HP (Cytiva) ion exchange chromatography. Affinity tags were removed by incubation with TEV protease and cleaved samples were purified by HiTrap Heparin (Cytiva) affinity, reverse binding to amylose (NEB) and size exclusion chromatography (HiLoad^TM^ 16/600 Superdex 200, Cytiva) in 50 mM Tris pH 8.0, 150 mM NaCl, 10% glycerol, 2 mM DTT. Protein samples were concentrated using Amicon® Ultra centrifugal filters (Millipore), and were stored at −80°C following flash-freezing in liquid nitrogen. Protein samples were analysed by SDS-PAGE with Coomassie staining, and concentrations were determined by UV spectroscopy using a Cary 60 UV spectrophotometer (Agilent) with extinction coefficients and molecular weights calculated by ProtParam (http://web.expasy.org/protparam/).

### Size-exclusion chromatography multi-angle light scattering (SEC-MALS)

The absolute molecular masses of human and *At* HEI10 constructs were determined by size-exclusion chromatography multi-angle light scattering (SEC-MALS). Protein samples at 5-10 mg/ml were loaded onto a Superdex™ 200 Increase 10/300 GL size exclusion chromatography column (Cytiva) in 50 mM Tris pH 8.0, 150 mM NaCl, 10% glycerol, at 0.5 ml/min using an ÄKTA™ Pure (Cytiva). The column outlet was fed into a DAWN® HELEOS™ II MALS detector (Wyatt), followed by an Optilab® T-rEX™ differential refractometer (Wyatt). Light scattering and differential refractive index data were collected and analysed using ASTRA® 6 software (Wyatt). Molecular weights and estimated errors were calculated across eluted peaks by extrapolation from Zimm plots using a dn/dc value of 0.1850 ml/g. SEC-MALS data are presented as differential refractive index (dRI) profiles, with fitted molecular weights (M_W_) plotted across elution peaks.

### Circular dichroism (CD) spectroscopy

Far UV circular dichroism (CD) spectroscopy data were collected on a Jasco J-810 spectropolarimeter (Institute for Cell and Molecular Biosciences, Newcastle University). CD spectra were recorded in 10mM Na_2_HPO_4_/ NaH_2_PO_4_ pH 7.5, at protein concentrations between 0.1-0.5 mg/ml, using a 0.2 mm pathlength quartz cuvette (Hellma), at 0.2 nm intervals between 260 and 185 nm at 4°C. Spectra were averaged across nine accumulations, corrected for buffer signal, smoothed and converted to mean residue ellipticity ([θ]) (x1000 deg.cm^2^.dmol^-1^.residue^-1^). Deconvolution was performed using the CDSSTR algorithm of the Dichroweb server (http://dichroweb.cryst.bbk.ac.uk) (Sreerama and Woody 2000; Whitmore and Wallace 2008). CD thermal denaturation was performed in 20 mM Tris pH 8.0, 150 mM KCl, 2 mM DTT, at protein concentrations between 0.1-0.4 mg/ml, using a 1 mm pathlength quartz cuvette (Hellma). Data were recorded at 222 nm, between 5°C and 95°C, at 0.5°C intervals with ramping rate of 2°C per minute, and were converted to mean residue ellipticity ([θ_222_]) and plotted as % unfolded ([θ]_222,x_-[θ]_222,5_)/([θ]_222,95_-[θ]_222,5_). Melting temperatures (Tm) were estimated as the points at which samples are 50% unfolded.

### Size-exclusion chromatography small-angle X-ray scattering (SEC-SAXS)

SEC-SAXS experiments were performed at beamline B21 of the Diamond Light Source synchrotron facility (Oxfordshire, UK). Protein samples at concentrations >5 mg/ml were loaded onto a Superdex™ 200 Increase 10/300 GL size exclusion chromatography column (Cytiva) in 20 mM Tris pH 8.0, 150 mM KCl at 0.5 ml/min using an Agilent 1200 HPLC system. The column outlet was fed into the experimental cell, and SAXS data were recorded at 12.4 keV, detector distance 4.014 m, in 3.0 s frames. Data were subtracted and averaged, and analysed for Guinier region *Rg* using ScÅtter 3.0, and *P(r)* distributions were fitted using *PRIMUS* (P.V.Konarev 2003). Crystal structures and models were fitted to experimental data using *FoXS* (Schneidman-Duhovny et al. 2016), and flexible termini were modelled and fitted to experimental data using *CORAL* (Petoukhov et al. 2012).

### Spectrophotometric determination of zinc content

The presence of zinc in protein samples was determined through a spectrophotometric method using the metallochromic indicator 4-(2-pyridylazo) resorcinol (PAR) (Sabel, Shepherd, and Siemann 2009). Protein samples at 20 µM were digested with 0.6 μg/μl proteinase K (NEB) at 60°C for 1 hr. Following centrifugation, 10 μl from the supernatant of each protein digestion was added to 80 μl of 50 μM 4-(2-pyridylazo)-resorcinol (PAR) in 20 mM Tris pH 8.0, 150 mM KCl, incubated for 5 min at room temperature, and UV absorbance spectra were recorded between 600 and 300 nm (Varian Cary 60 spectrophotometer). Zinc concentrations were estimated from the ratio between absorbance at 492 and 414 nm, plotted on a line of best fit obtained from analysis of 0–100 μM ZnOAc standards.

### Crystallisation and structure solution of HEI10 core

Protein crystals of HEI10 core were obtained through vapour diffusion in hanging drops, by mixing 1 µl of protein at 10 mg/ml with 1 µl of crystallisation solution (0.1 M HEPES sodium salt pH 7.5 30% (w/v) MPD; 5% (w/v) PEG 4000) and equilibrating at 20°C. Crystal quality was improved by micro-seeding, using a 1:1000 seed stock solution in the above crystallisation condition. Crystals were cryo-cooled in liquid nitrogen. X-ray diffraction data were collected at 1.27806 Å, 100 K, as 3600 consecutive 0.10° frames of 0.050 s exposure on an Eiger2 XE 16M detector at beamline I03 of the Diamond Light Source synchrotron facility (Oxfordshire, UK). Data were processed using AutoPROC (Vonrhein et al. 2011), in which indexing, integration, scaling and merging were performed by XDS (Kabsch 2010) and Aimless (Evans 2011), and anisotropic correction with a local I/σ(I) cut-off of 1.2 was performed by STARANISO (Tickle et al. 2018). Crystals belong to monoclinic spacegroup P2_1_ (cell dimensions a = 53.20 Å, b = 73.86 Å, c = 124.03 Å, α = 90°, β = 99.28°, γ = 90°), with one HEI10 tetramer in the asymmetric unit. Structure solution was achieved through molecular replacement by Phaser (McCoy et al. 2007) using an *AlphaFold 2* model of the centre of the HEI10 tetramer (amino-acids 1-122) as a search model (Evans et al. 2021; Jumper et al. 2021). Model building was performed through iterative re-building by *PHENIX Autobuild* (Adams et al. 2010) and manual building in *Coot* (Emsley et al. 2010). The structure was refined using PHENIX refine (Adams et al. 2010), with isotropic atomic displacement parameters with eleven TLS (translation–libration–screw rotation) groups. The structure was refined against anisotropy-corrected data with resolution limits between 2.4 Å and 5.7 Å, to R and R_free_ values of 0.2498 and 0.2891, respectively, with 98.30% of residues within the favoured regions of the Ramachandran plot (0 outliers), clashscore of 6.03 and overall MolProbity score of 1.33 (Chen et al. 2010).

### Structural modelling

Models of human HEI10 1-122 were generated using *AlphaFold 2* (Jumper et al. 2021). Models of *Arabidopsis thaliana* HEI10, bound to two zinc ions per chain, were generated by *AlphaFold 3* using the *AlphaFold* Server (Abramson et al. 2024). The pLDDT and PAE plots were generated using https://github.com/Ash100/Biopython/blob/main/AF3_Results_Visualization.ipynb. The HEI10 core crystal structure (in which residues up to 184 were visible) was extended up to the C-terminal boundary of residue 196 for fitting to SAXS data and interpretation of fibrous contacts within the crystal lattice. To achieve this, a coiled-coil model of residues 171-196 was generated by *CCBuilder 2.0* (Wood and Woolfson 2018), and was docked onto each end of the crystal structure and connected using *Coot* (Emsley et al. 2010).

### Electron microscopy

Electron microscopy was performed using a JEOL JEM-1400 Plus transmission electron microscope equipped with a Gatan OneView camera (Structural Biology Core of the Discovery Research Platform for Hidden Cell Biology, University of Edinburgh). Carbon-coated copper grids, 300 mesh, were glow discharged using a PELCO easiGlow glow discharge system at 25 mA for 60 secs. Protein samples at 0.005–3 mg ml^−1^ were applied to the glow discharged grids, following by negative staining with 2% (weight to volume) uranyl acetate.

### HEI10 transient expression and co-immunofluorescence analysis in COS7 cells

Different sequences of human HEI10 and *At* HEI10 were cloned into the pcDNA™3.1 expression vector (Invitrogen®) as fusion proteins with an N-terminal 3xFLAG tag (DYKDDDDK) for constitutive expression in mammalian cells. COS-7 cells (ATCC CRL-1651) were seeded at 1:5 dilution onto poly-L-lysine treated 18x18 mm HiQA high-performance coverslips (CellPath®) and grown in high-glucose Dulbecco′s Modified Eagle′s Medium (Sigma-Aldrich®) supplemented with 10% FBS and 1% Pen Strep (Gibco®) at 37°C and 5% CO_2_. Once the cells reached 90% confluence these were transfected with the expression vector using Lipofectamine 3000 Transfection Reagent (Invitrogen®) following the instructions of the manufacturer. Constitutive expression of the HEI10 constructs was allowed for 24h prior to the preparation of the cells for immunofluorescence.

After 24h the transfected COS-7 cells were washed once with 1xPBS buffer (137 mM NaCl, 2.7 mM KCl, 10 mM Na_2_HPO_4_, and 1.8 mM K_2_HPO_4_; pH 7.4) and subsequently fixed with cold 100% methanol at −20°C for 20 minutes. Once the fixation was completed, the coverslips were left to air dry for 5 minutes. The coverslips were then washed three times for 5 min in PBS-T buffer (1xPBS, 0.1% Triton X-100) after which the cells were incubated in blocking buffer (3% Bovine Serum Albumin in PBS-T buffer) for 30 min at 37°C in a moist chamber. Subsequently, the cells were incubated overnight at 4°C with the primary antibody (mouse anti-FLAG® M2 monoclonal IgG, Sigma-Aldrich®; 1:250 dilution in blocking buffer). The coverslips were then washed three times for 5 minutes with PBS-T and incubated with the secondary antibody (Goat anti-Mouse IgG Alexa Fluor® 594, Invitrogen®; 1:250 dilution in blocking buffer) for 90 min at RT. Next, the coverslips were washed three times for 5 minutes with PBS-T and stained with Hoechst 33342 (10 µg/ml in 1xPBS) for 10 minutes at RT and subsequently washed once with PBS-T buffer. Finally, the coverslips were mounted on a glass slide with 10 µl of Vectashield® anti-fading mounting medium (Vector Laboratories®) and stored at −20°C.

Widefield imaging of immunofluorescence slides was conducted using a Nikon® Ti2 Widefield microscope equipped with a Photometrics Prime 95B camera (Light Microscopy Core of the Discovery Research Platform for Hidden Cell Biology, University of Edinburgh). Super-resolution fluorescence microscopy imaging was performed using a Nikon® Ti2 microscope equipped with a Nikon® CSU-W1 SoRa system and a Photometrics® Prime 95B camera (Advanced Imaging Resource, University of Edinburgh). Fluorescence microscopy images were analysed and processed using the NIS Image Viewer (Nikon®) and the Fiji / ImageJ software. In addition, super-resolution fluorescence images were deconvolved prior to their analysis using the deconvolution tool from the NIS Image Analysis software (Nikon®). Image panels were assembled using Adobe Illustrator and Adobe Photoshop 2020 (Adobe®).

### HEI10 transient expression and co-immunofluorescence analysis in U2OS cells

U2OS cells were seeded at 2 × 10⁵ cells per well in 6-well plates on sterile glass coverslips in DMEM (Thermo Fisher Scientific, 41966029) supplemented with 5% fetal bovine serum and penicillin–streptomycin (10,000 U/mL, Thermo Fisher Scientific) and incubated at 37 °C with 5% CO₂. The following day, cells were transiently transfected with 2 µg plasmid DNA using 4 µL Lipofectamine 2000 (Thermo Fisher Scientific, 11668027) in Opti-MEM reduced-serum medium (Thermo Fisher Scientific, 15392402). The transfection mixture was added dropwise to cells.

At 24 h post-transfection, for DNA damage experiments, cells were treated with 1 µM gemcitabine or DMSO vehicle control for 24 h. At 48 h post-transfection, cells were washed once with ice-cold PBS and fixed in 100% ice-cold methanol at −20 °C for 10 min. After PBS rinsing, cells were incubated with primary antibodies in PBS for 1 h at room temperature in a humidified dark chamber. The following primary antibodies were used: anti-FLAG (mouse, 1:200, Sigma-Aldrich, F3165), anti-FLAG (rabbit, 1:200, Invitrogen, 740001), anti-PML (mouse, 1:500, Abcam, ab96051), anti-coilin (mouse, 1:400, Abcam, ab11822), anti-pericentrin (mouse, 1:200, Abcam, ab28144), anti-RAD51 (mouse, 1:100, Abcam, ab88572), anti-γH2AX (mouse, 1:400, Abcam, ab26350), and anti-PCM1 (rabbit, 1:100, Proteintech, 19856-1-AP).

Following three 5-min washes, cells were incubated with goat anti-mouse Alexa Fluor 488 (Invitrogen, A11001) and goat anti-rabbit Alexa Fluor 594 (Invitrogen, A11012) secondary antibodies (0.01 mg/mL) together with DAPI (0.02 mg/mL) for 1 h at room temperature in a humidified dark chamber. Cells were washed twice with PBS and once with Milli-Q water. Coverslips were air-dried for 15 min in the dark and mounted using ProLong Gold Antifade Mountant (Invitrogen, P36930).

Images were acquired using a Zeiss LSM880 confocal microscope. Colocalization analysis was performed in Fiji (ImageJ) using channel-specific thresholds that were applied consistently across all images, followed by generation of binary masks for each channel and quantification of overlap as the proportion of channel A signal within channel B (A over B) and channel B signal within channel A (B over A).

### Protein sequence and structure analysis

HEI10 sequences were aligned and visualised using MUSCLE (Madeira et al. 2019) and Jalview (Waterhouse et al. 2009). Molecular structure images were generated using the *PyMOL* Molecular Graphics System, Version 3.0 Schrödinger, LLC.

### Accession codes and data availability

Crystallographic structure factors and atomic co-ordinates have been deposited in the Protein Data Bank (PDB) under accession number 9GES, and raw diffraction data have been uploaded to https://proteindiffraction.org/.

## Acknowledgements

We thank Diamond Light Source and the staff of beamlines I03 and B21 (proposals mx18598, sm15836, sm25084 and mx32705). We thank A. Baslé and H. Waller for assistance with X-ray crystallographic and CD data collection; the University of Edinburgh Advanced Imaging Resource (AIR) for their assistance in the collection of super-resolution fluorescence microscopy data; and Nitzen Shabek and Jawdat Al-Bassam (UC Davis) for vectors and constructs. This work was supported by funding for the Wellcome Discovery Research Platform for Hidden Cell Biology (226791) and we gratefully acknowledge support from the Light Microscopy and Structural Biology cores; Eunice Kennedy Shriver National Institute of Child Health and Human Development award 5R01HD109322 to N.H.; Medical Research Council (MRC) (MR/X00855X/1), European Molecular Biology Organization Young Investigator Network Installation Grant and NCN Sonata Bis grant no. 2023/50/E/NZ3/00281 to U.L.M.; Medical Research Council University Unit award (MC_UU_00035/3) to I.R.R.. O.R.D. is a Wellcome Senior Research Fellow (Grant Number 219413/Z/19/Z). M.E.W. is supported by University of Liverpool Doctoral Scholarship. N.H. is an Investigator with the Howard Hughes Medical Institute.

## Author contributions

A.E.M., C.E.S. and J.M.D. purified and performed biophysical experiments on human HEI10, and crystallised the HEI10 core. D.S.K purified human HEI10, performed ubiquitylation assay and prepared figures. J.N. and S.D. purified and performed biophysical experiments on *Arabidopsis thaliana* HEI10. M.P.P. and M.G. performed EM on HEI10 full-length. M.P.P. and M.E.W. performed cellular over-expression experiments. O.R.D. solved the HEI10 core crystal structure. O.R.D., U.L.M. I.R.A. and N.H. analysed data, designed and interpreted experiments and wrote the manuscript.

## Declaration of interests

The authors declare no competing interests.

---

**Supplementary Figure 1.**
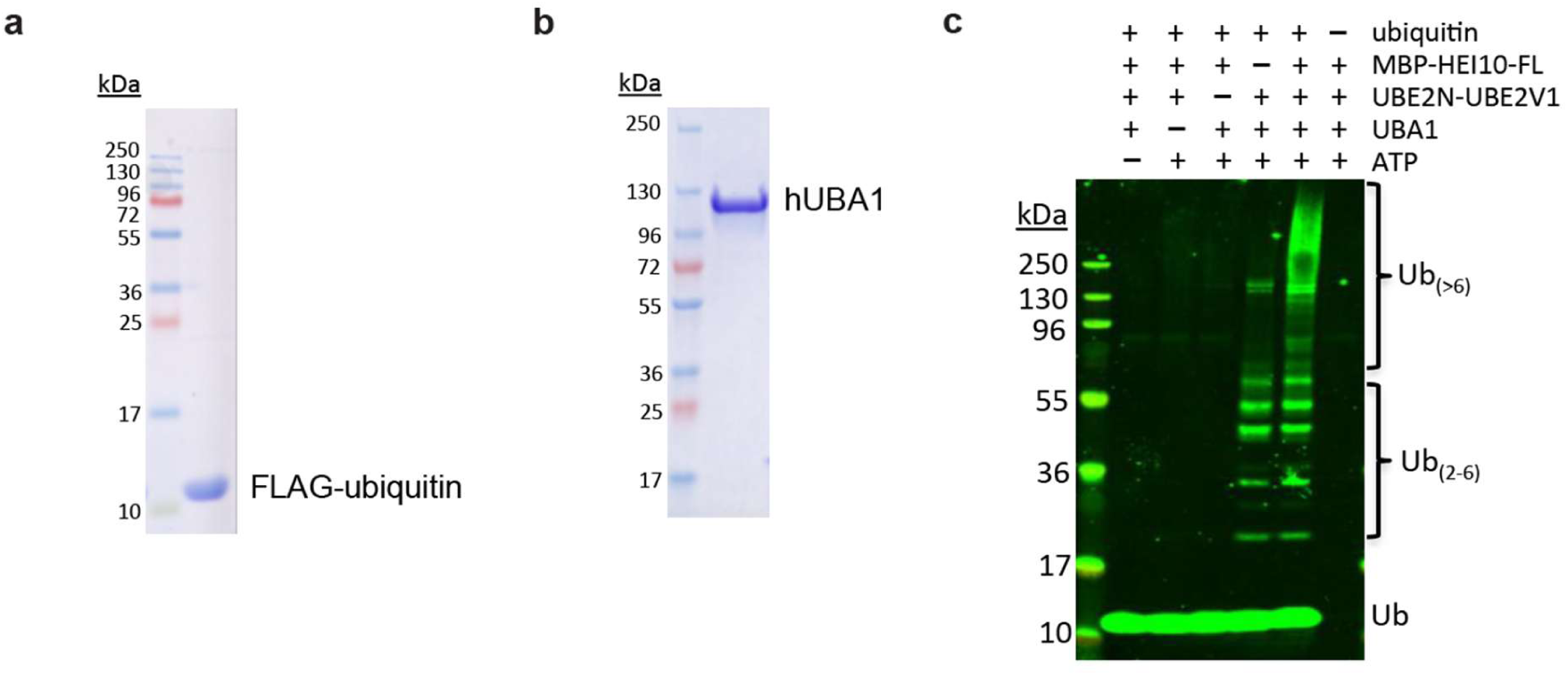
HEI10 catalyses ubiquitin chain formation *in vitro*. (**a,b**) SDS-PAGE analysis of purified FLAG-ubiquitin and human UBA1 (Boston Biochem) used in ubiquitylation reactions. (**c**) Western blot image of control ubiquitylation reactions showing that high-molecular weight Ub_(>6)_ chains require all reaction components.

**Supplementary Figure 2.**
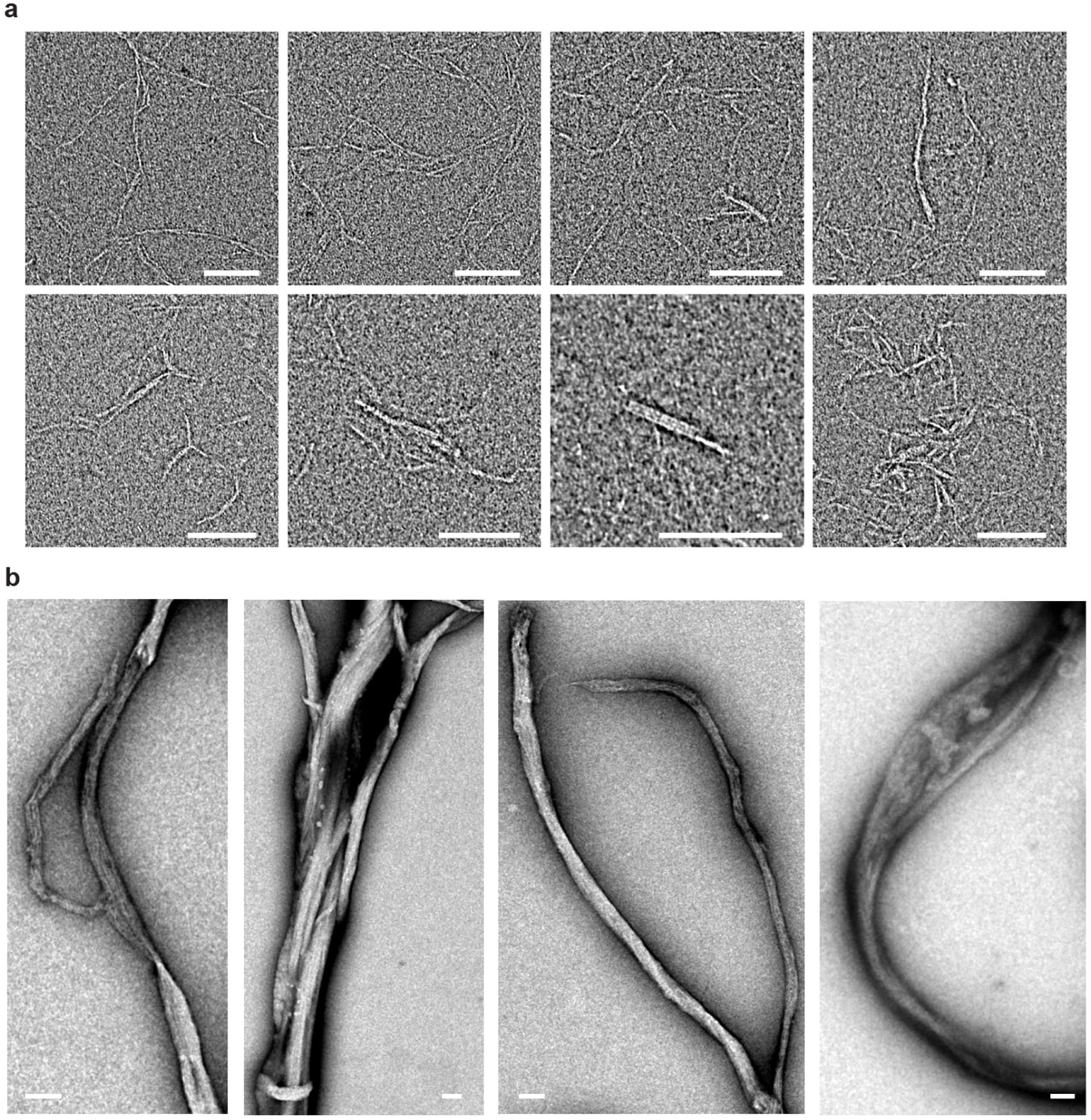
Fibrous self-assembly of HEI10 *in vitro*. Additional electron micrographs of HEI10 (full-length) pellets following TEV-cleavage from its N-terminal MBP tag, showing (**a**) internal fibrillar structure within HEI10 fibres and (**b**) large fibrous assemblies, supporting data shown in Figure 1e,f. Scale bars, 100 nm.

**Supplementary Figure 3.**
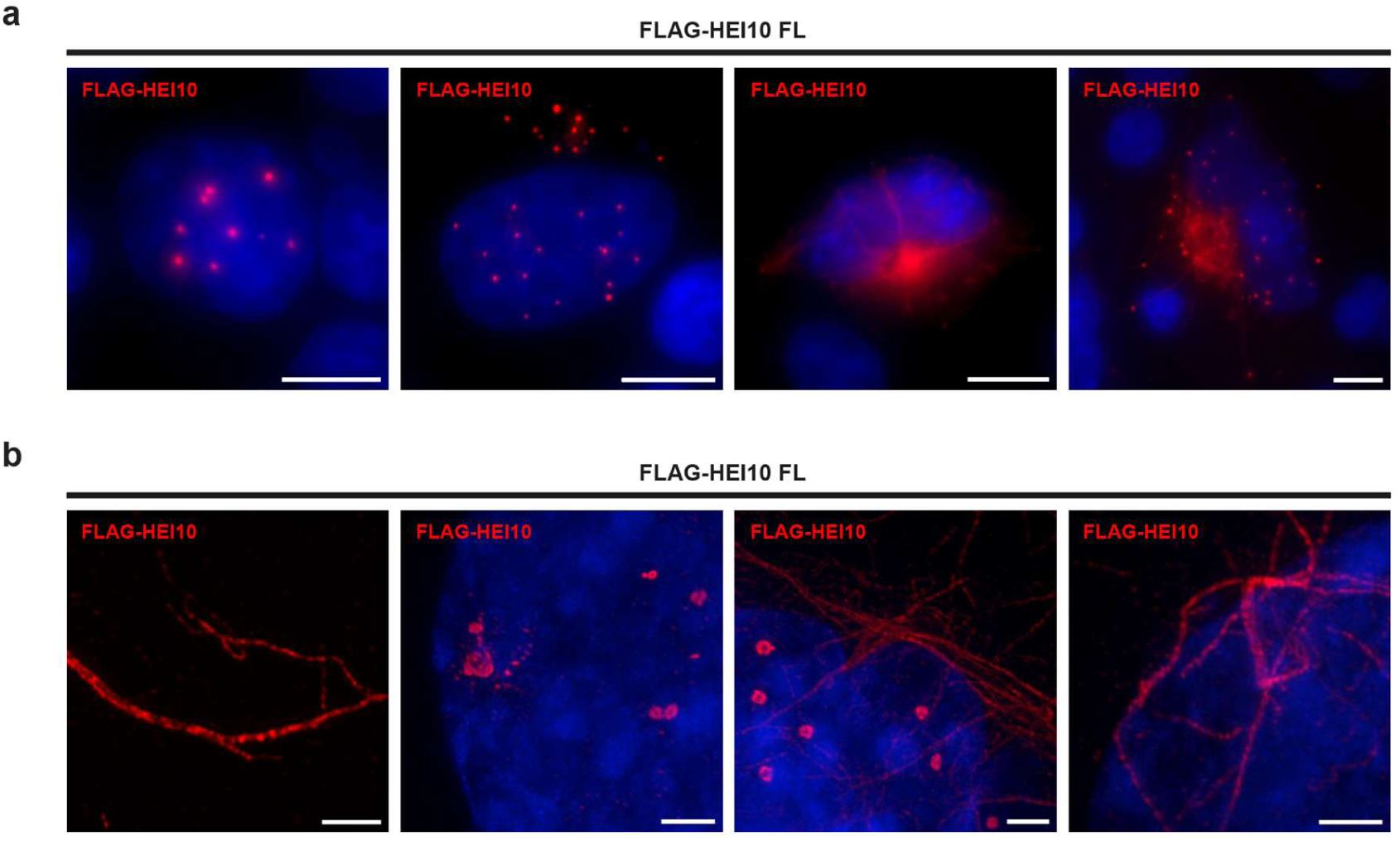
HEI10 forms cytoplasmic fibres and nuclear/cytoplasmic foci upon cellular expression. (**a,b**) Additional replicates of Flag-HEI10 over-expression in COS7 cells, with staining for Flag-HEI10 (red) and Hoechst 33342 (blue), supporting data shown in Figure 2a,b. (**a**) Wide-field imaging, showing that Flag-HEI10 forms cytoplasmic fibres and nuclear foci. Scale bars: 10 μm. (**b**) SoRa super-resolution imaging of cytoplasmic fibres (left) and nuclear foci (right). Scale bars: 2 μm.

**Supplementary Figure 4.**
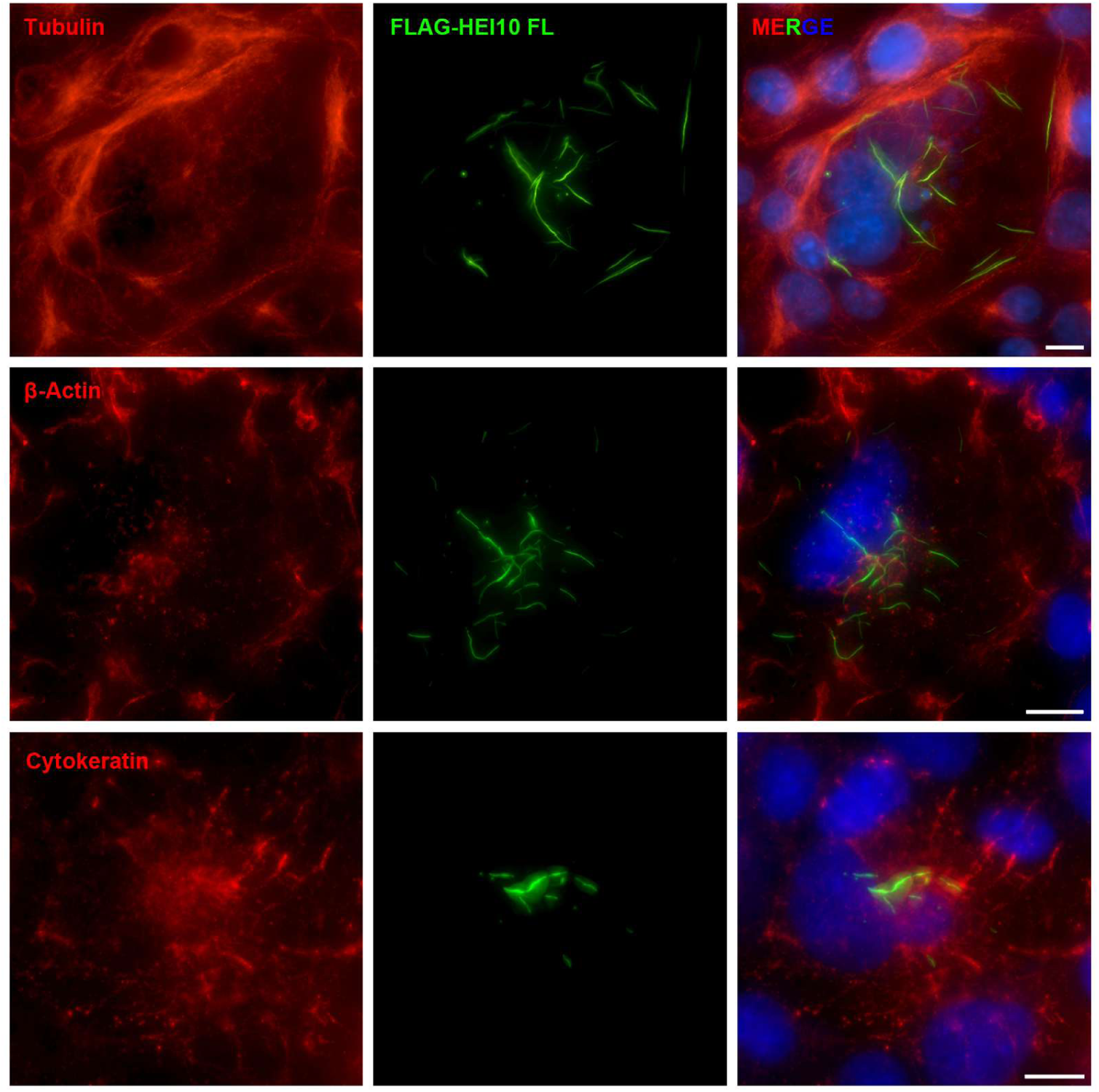
Fibrous assembly of HEI10 upon expression in somatic cells. Transient over-expression of Flag-HEI10 in COS7 cells, analysed by wide-field imaging, co-stained for Flag-HEI10 (red), DAPI, and cytoskeletal components tubulin, β-actin and cytokeratin. Scale bars, 10 μm.

**Supplementary Figure 5.**
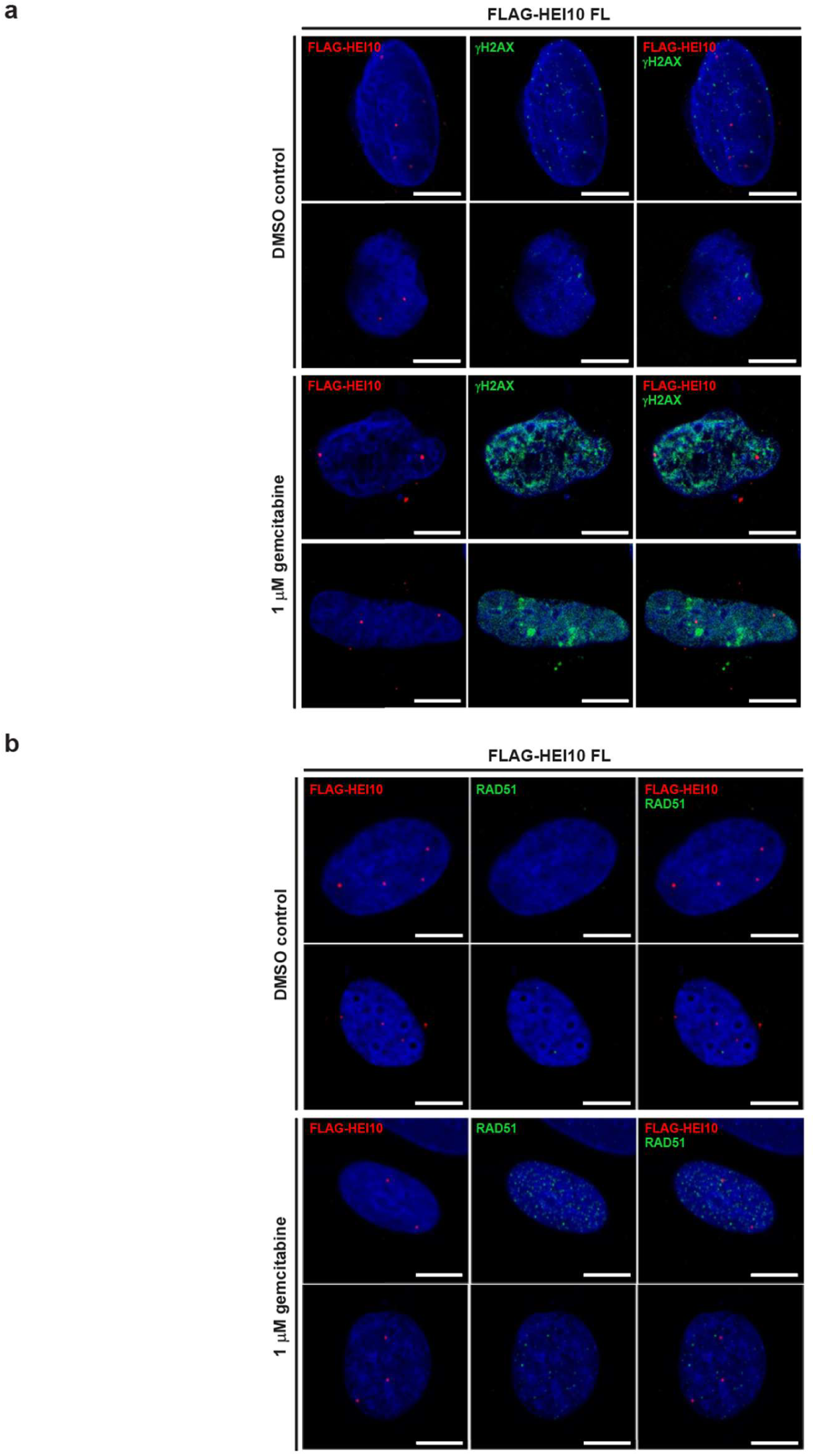
HEI10 foci do not co-localise with DNA double-strand breaks. (**a,b**) Additional replicates of Flag-HEI10 over-expression in U2OS cells, showing co-staining of Flag-HEI10 (red), with (**a**) γH2AX (green) and (**b**) RAD51 (green), alongside DAPI (blue), in cells treated with a DMSO control and 1 μM gemcitabine, supporting data shown in Figure 2c,d. Scale bars, 10 μm.

**Supplementary Figure 6.**
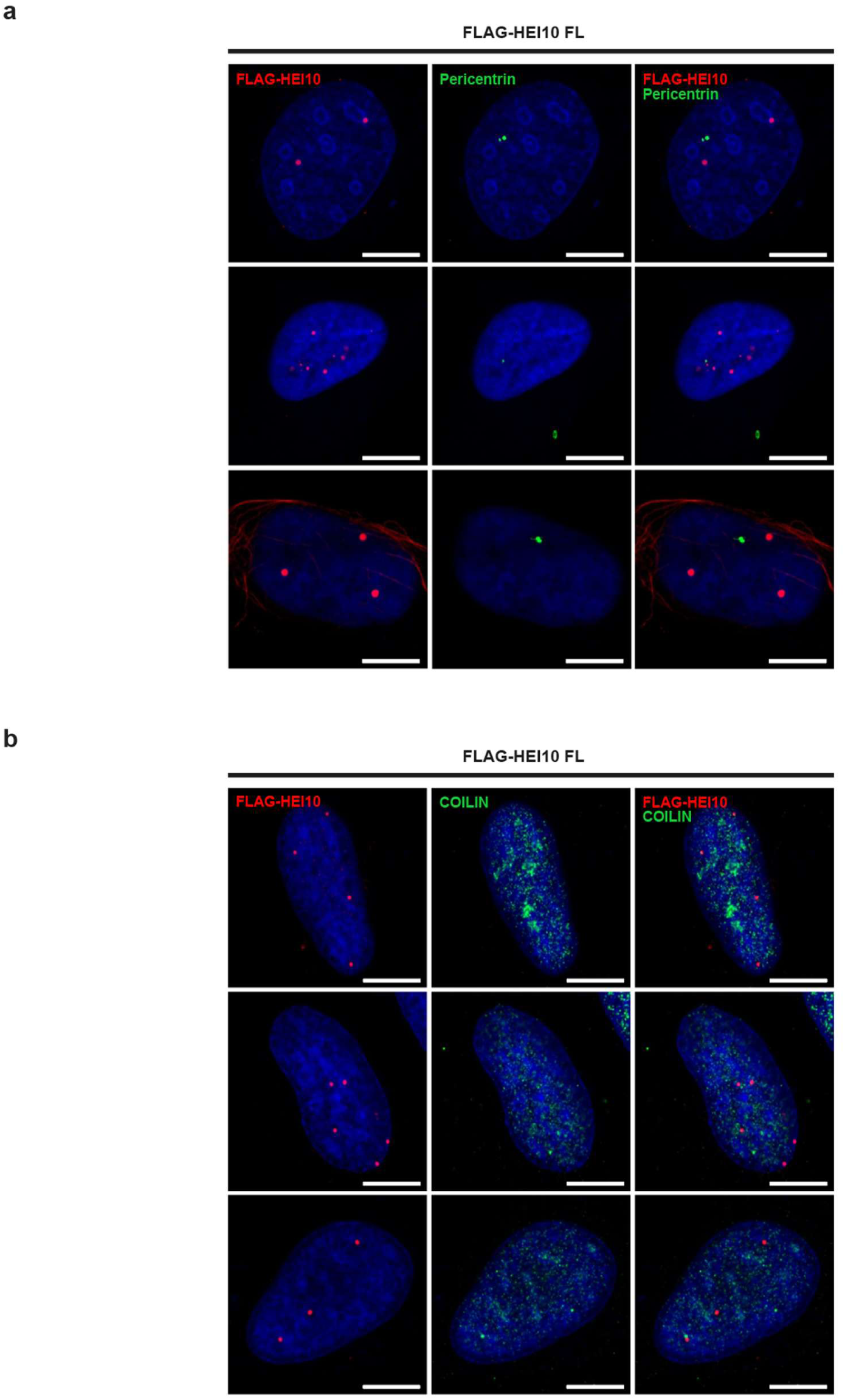
HEI10 foci do not co-localise with centrosomes or Cajal bodies. (**a,b**) Flag-HEI10 over-expression in U2OS cells. Co-staining of Flag-HEI10 (red), DAPI (blue) and (**a**) pericentrin (green) and (**b**) COILIN (green). Scale bars, 10 μm.

**Supplementary Figure 7.**
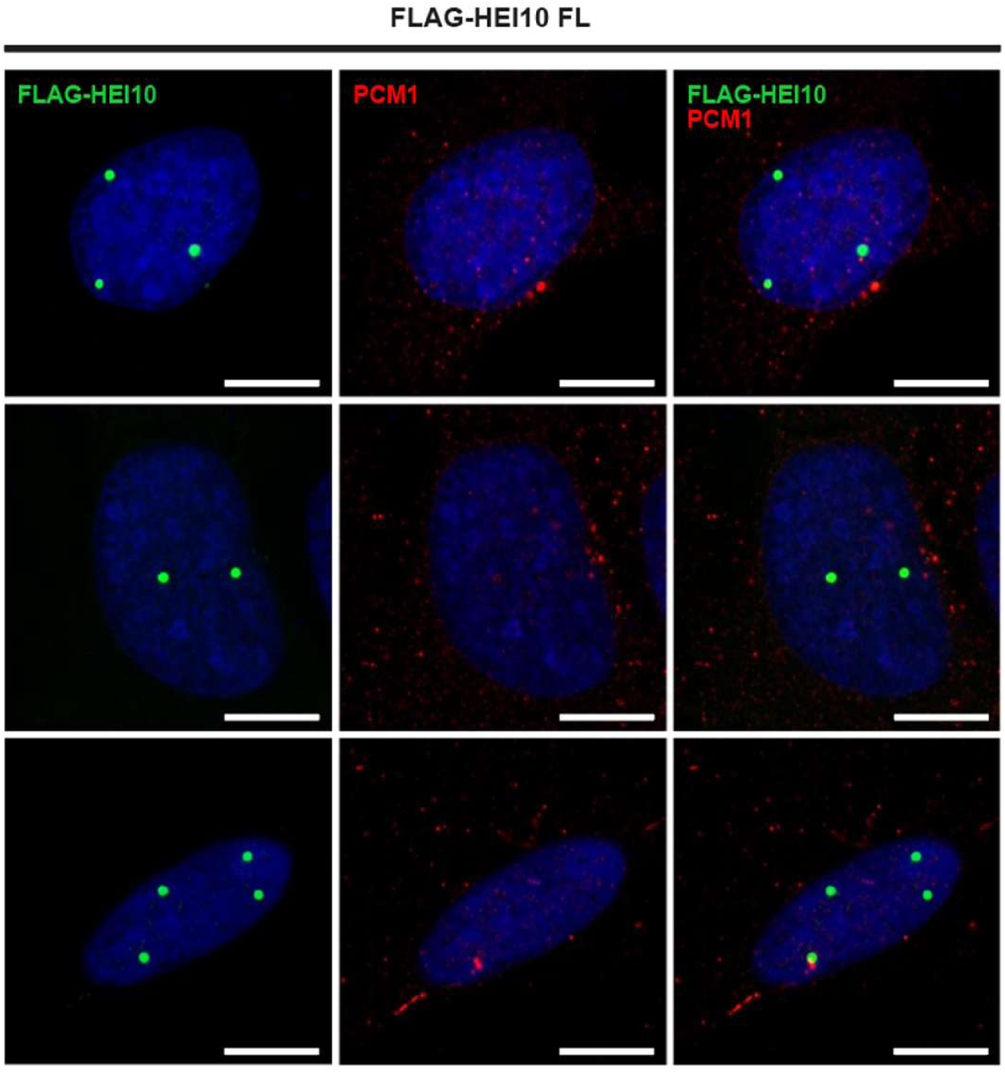
HEI10 foci do not co-localise with peri-centriolar material. Flag-HEI10 over-expression in U2OS cells. Co-staining of Flag-HEI10 (red), DAPI (blue) and PCM1 (green). Scale bars, 10 μm.

**Supplementary Figure 8.**
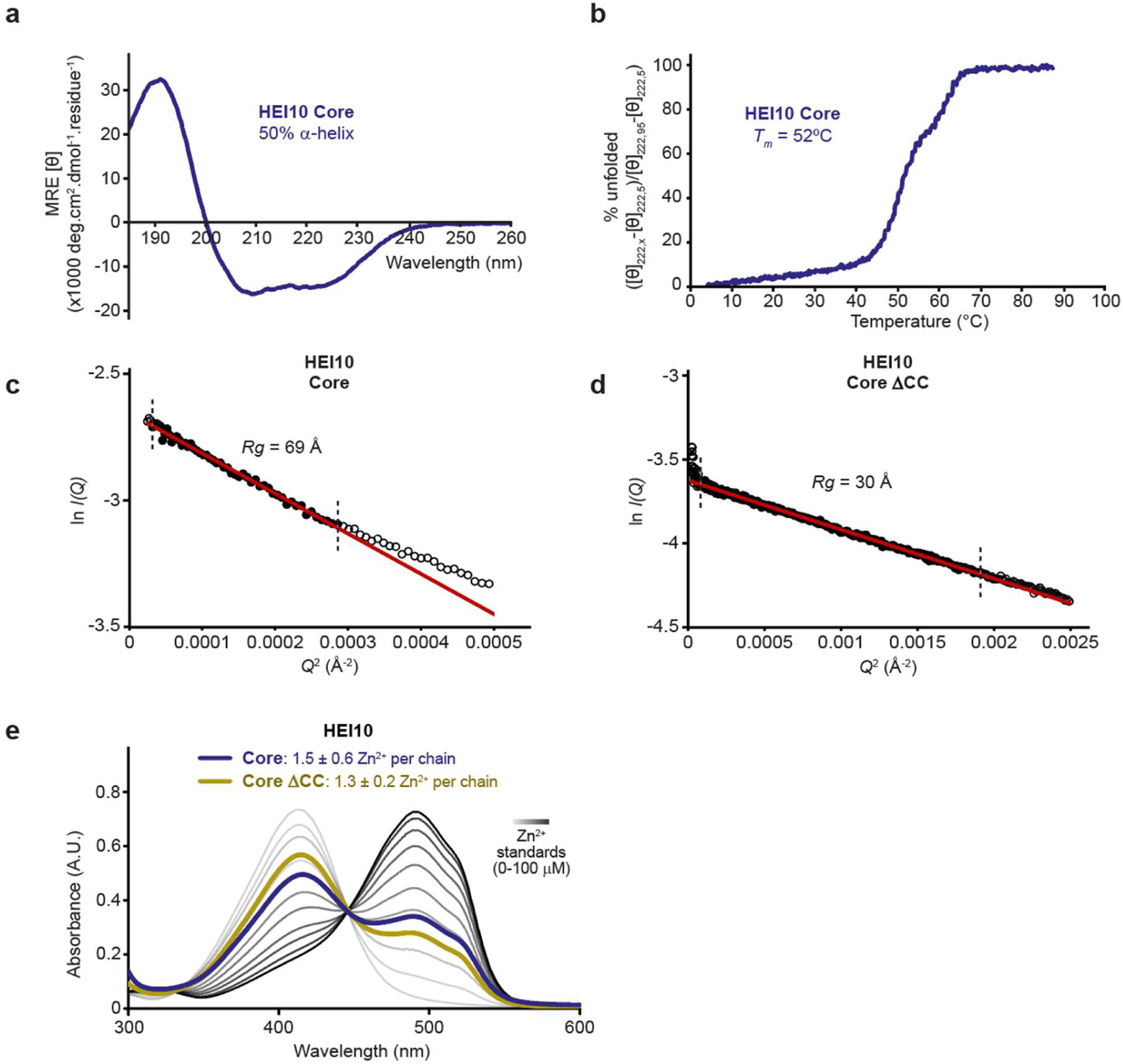
HEI10 core is a stable homo-tetramer in solution that corresponds to its crystal structure. (**a**,**b**) Circular dichroism (CD) analysis of HEI10 core. (**a**) Far UV CD spectra recorded between 260 nm and 185 nm in mean residue ellipticity, MRE ([θ]) (x10^3^ deg.cm^2^.dmol^−1^.residue^−1^). Data were deconvoluted using the CDSSTR algorithm, determining a helical content of 50%. (**b**) CD thermal denaturation of HEI10 core, recording the CD helical signature at 222 nm between 5°C and 95°C as % unfolded, indicating a melting temperature of 52°C. (**c**,**d**) SAXS Guinier analysis to determine the radius of gyration (*Rg*); linear fits are shown in red, with the fitted data range highlighted in black and demarcated by dashed lines. The *Q*.*Rg* values were < 1.3 and *Rg* was calculated for (**c**) HEI10 core and (**d**) HEI10 ΔCC as 69 Å and 30 Å, respectively. Corresponding to SEC-SAXS data shown in Figure 3c,d. (**e**) Spectrophotometric determination of zinc content for HEI10 core (dark blue; 1.5 ± 0.6 Zn^2+^ per chain, n = 6) and HEI10 1-122 (yellow; 1.3 ± 0.2 Zn^2+^ per chain, n = 3), using metallochromic indicator PAR, with zinc standards shown in a gradient from light to dark grey (0-100 µM) and proteins analysed at 20 µM (per chain).

**Supplementary Figure 9.**
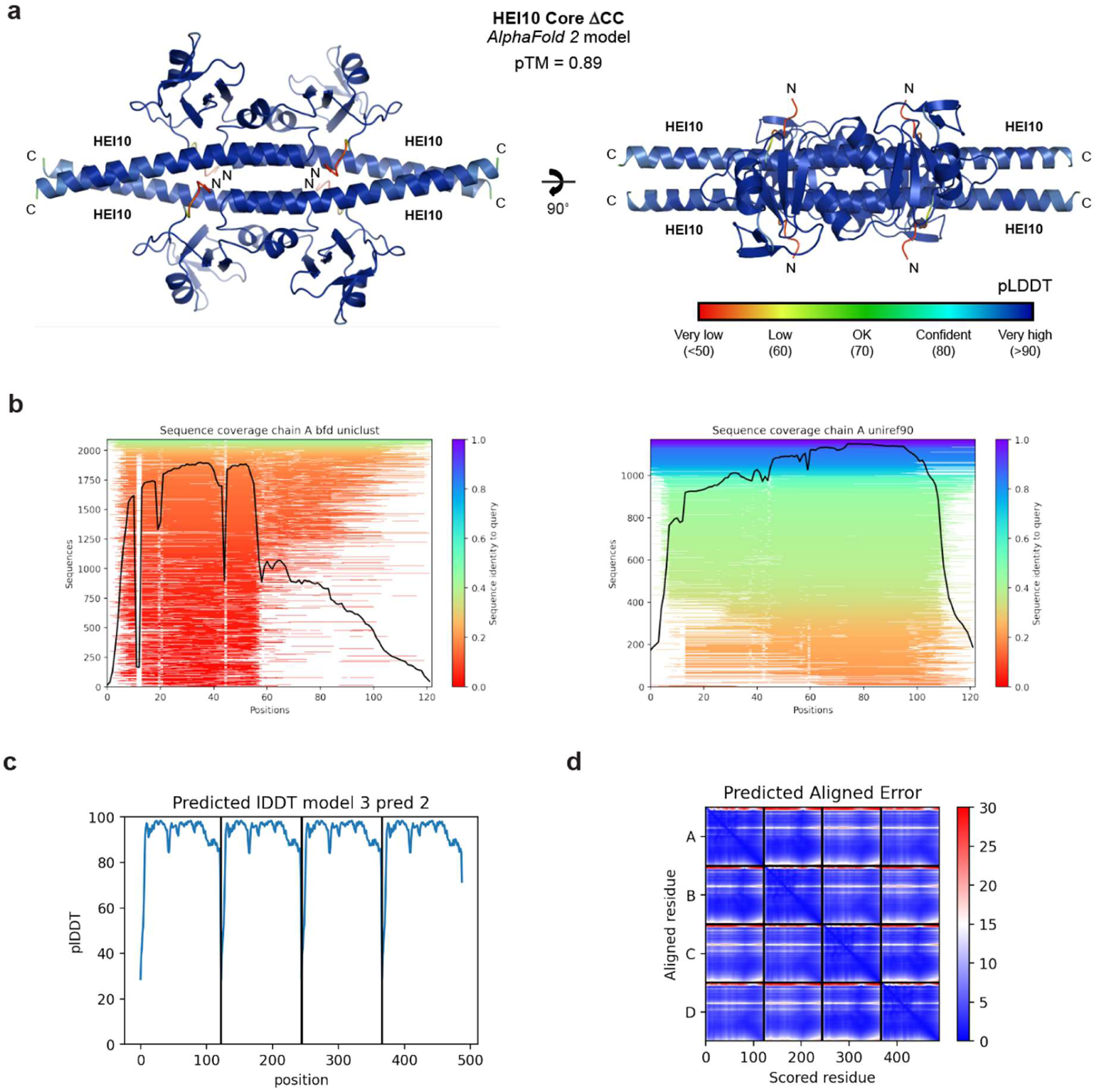
***AlphaFold 2* model of the HEI10 core ΔCC tetramer.** (**a**) *AlphaFold 2* model of the HEI10 core ΔCC tetramer coloured according to predicted LDDT (pLDDT) scores, between blue (>90) and red (<50). (**b**) Representations of the multiple sequence alignments generated and used by *AlphaFold 2*, showing the number of sequences and sequence identity against the position along the HEI10 sequence. (**c**) Predicted LDDT (pLDDT) scores shown for each amino-acid of the four HEI10 chains. (**e**) Predicted aligned error scores between each amino-acid of the four HEI10 chains, between blue (low error) and red (high error).

**Supplementary Figure 10.**
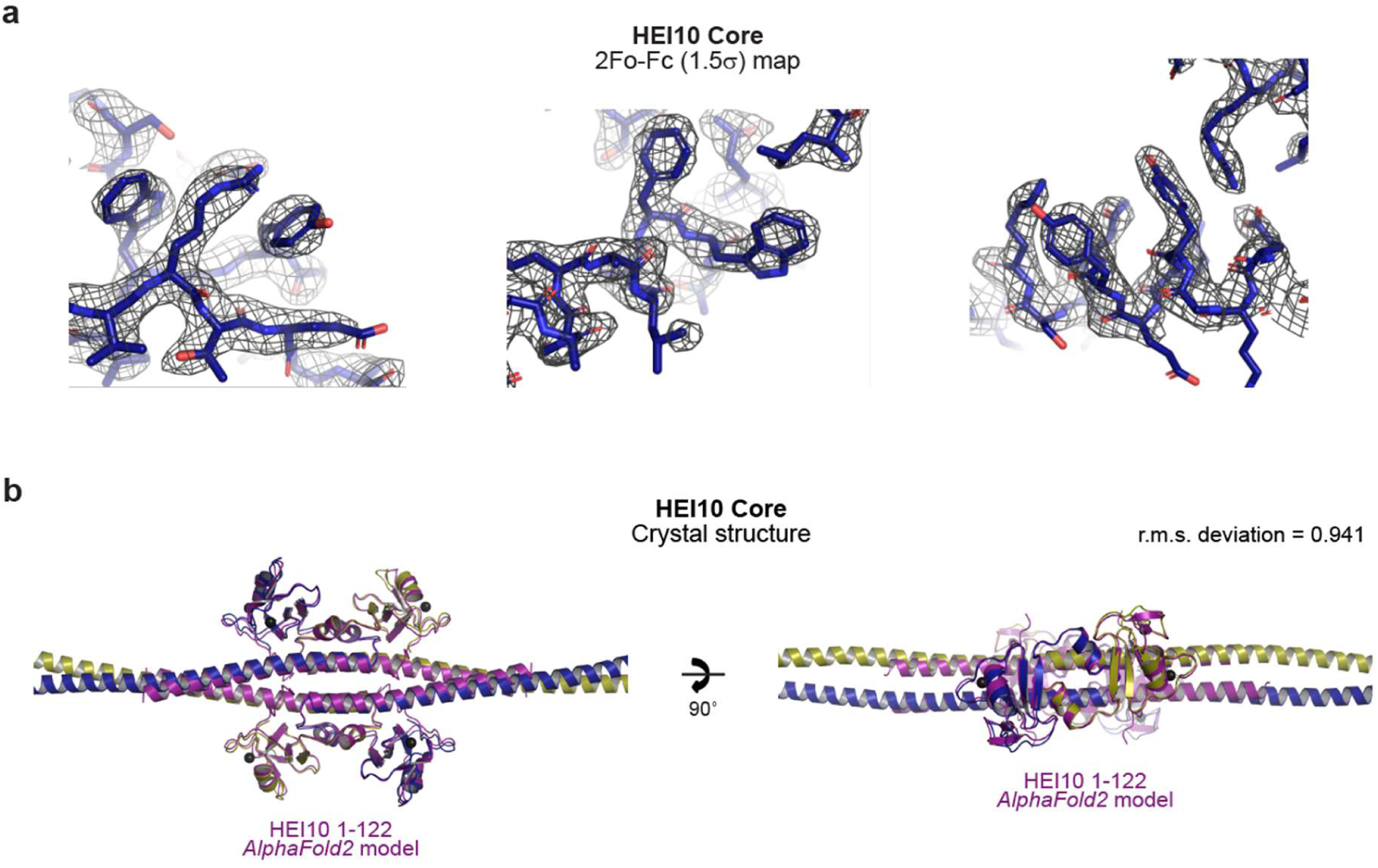
Crystal structure of HEI10 core. (**a**) 2Fo-Fc electron density map of the HEI10 core structure, contoured at 1.5σ, superimposed on the refined crystallographic model. (**b**) Superposition of the HEI10 core crystal structure (yellow and blue) with the HEI10 core ΔCC *AlphaFold 2* model (purple), showing an r.m.s. deviation of 0.941.

**Supplementary Figure 11.**
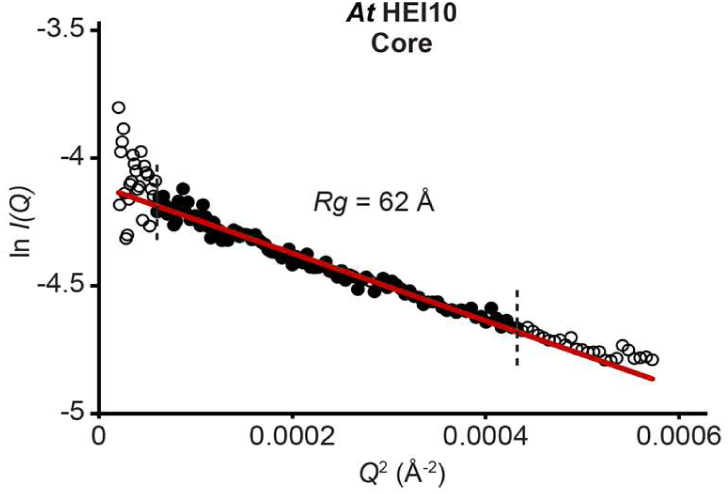
SEC-SAXS analysis of *At* HEI10 core. SAXS Guinier analysis to determine the radius of gyration (*Rg*); linear fits are shown in red, with the fitted data range highlighted in black and demarcated by dashed lines. The *Q*.*Rg* values were < 1.3 and *Rg* was calculated as 62 Å. Corresponding to SEC-SAXS data shown in Figure 6d,e.

**Supplementary Figure 12.**
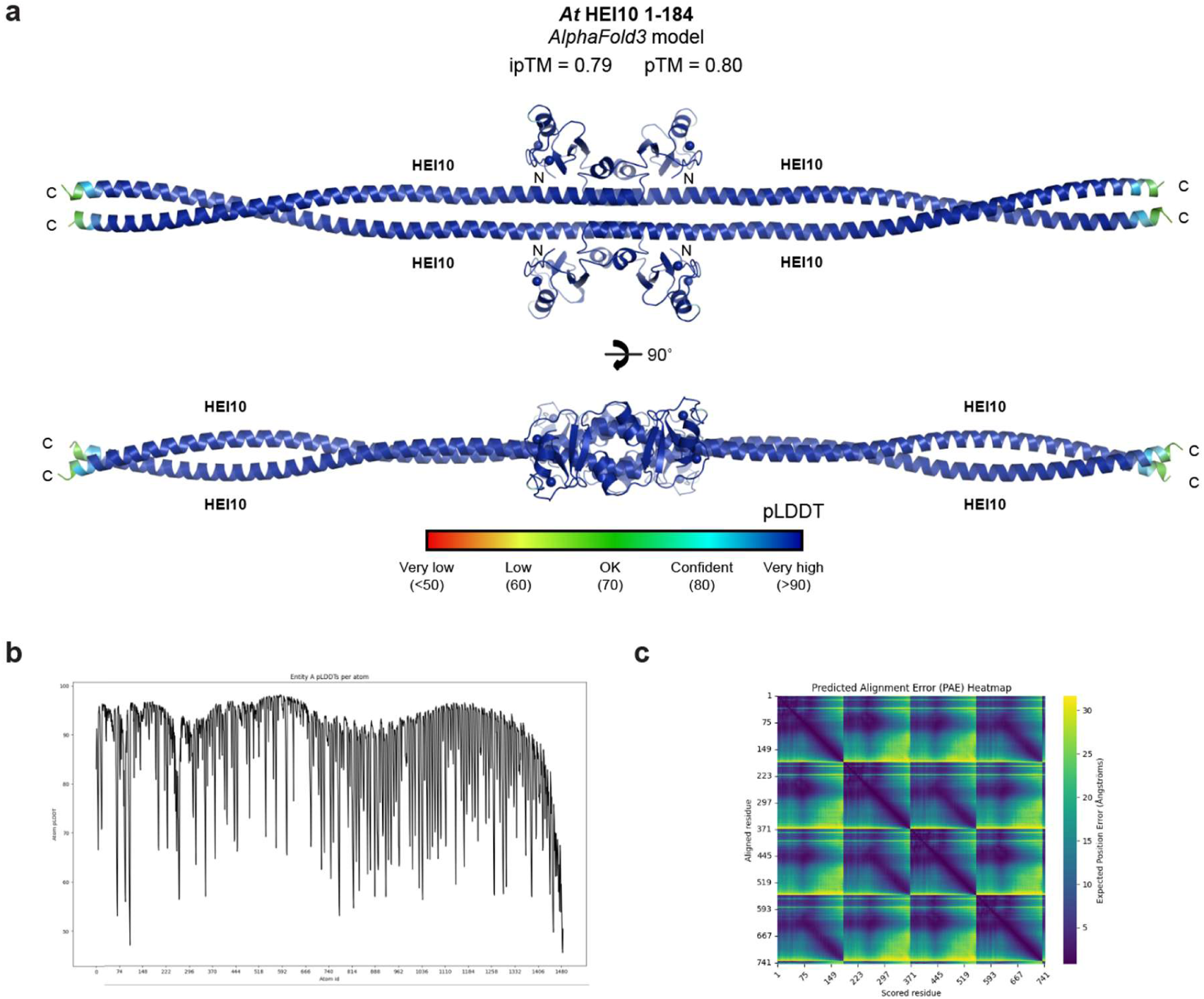
***AlphaFold 3* model of the *At* HEI10 core tetramer.** (**a**) *AlphaFold 3* model of the *At* HEI10 core tetramer coloured according to predicted LDDT (pLDDT) scores, between blue (>90) and red (<50). (**b**) Predicted LDDT (pLDDT) scores shown for each amino-acid of the four *At* HEI10 chains. (**c**) Predicted aligned error scores between each amino-acid of the four *At* HEI10 chains, between blue (low error) and yellow (high error).

**Supplementary Figure 13.**
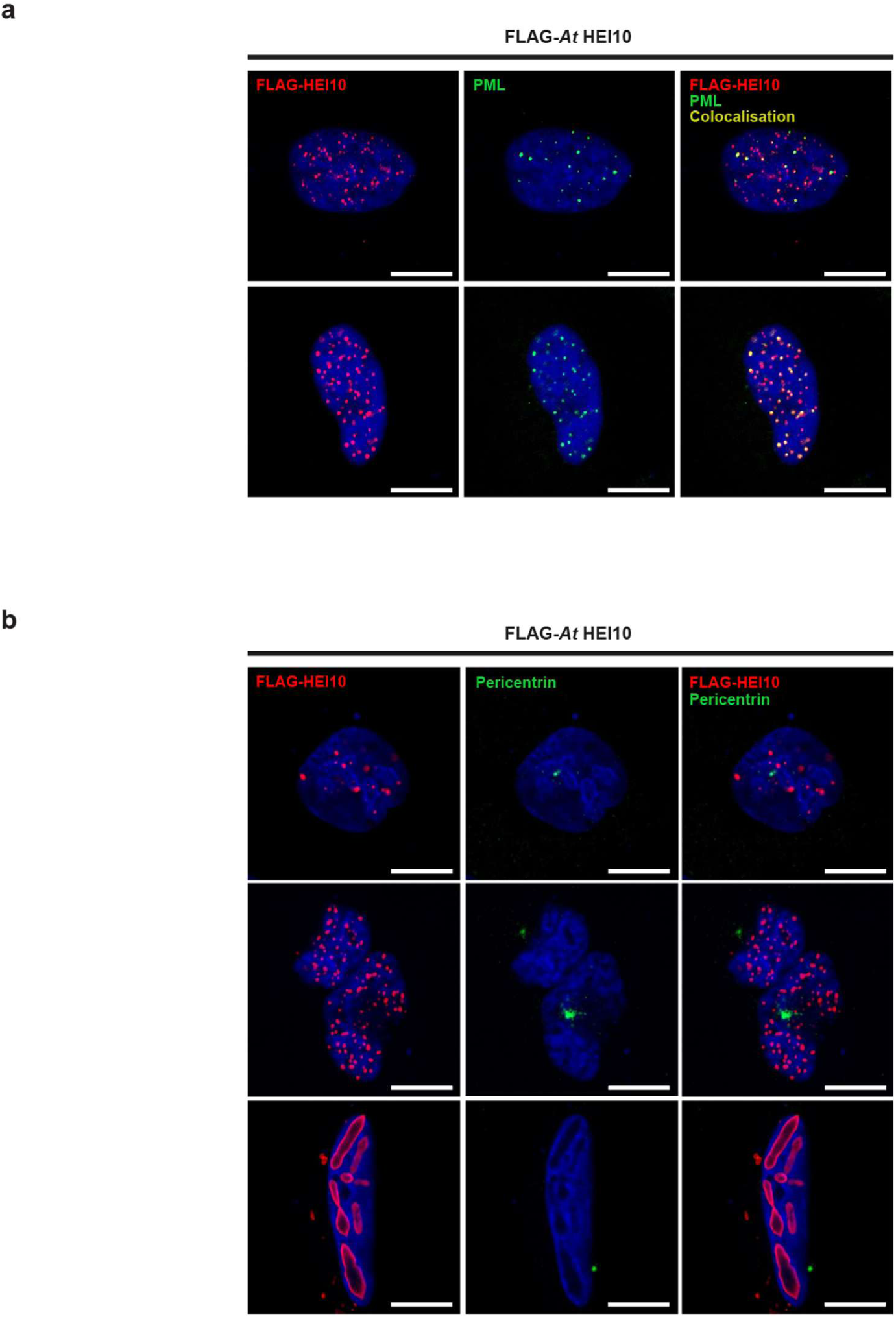
***At* HEI10 foci localise with PML bodies but not centrosomes.** (**a,b**) Flag-*At* HEI10 over-expression in U2OS cells. Co-staining of Flag-*At* HEI10 (red), DAPI (blue) and (**a**) PML (green) and (**b**) pericentrin (green), supporting data in Figure 6g. Scale bars, 10 μm.

**Supplementary Figure 14.**
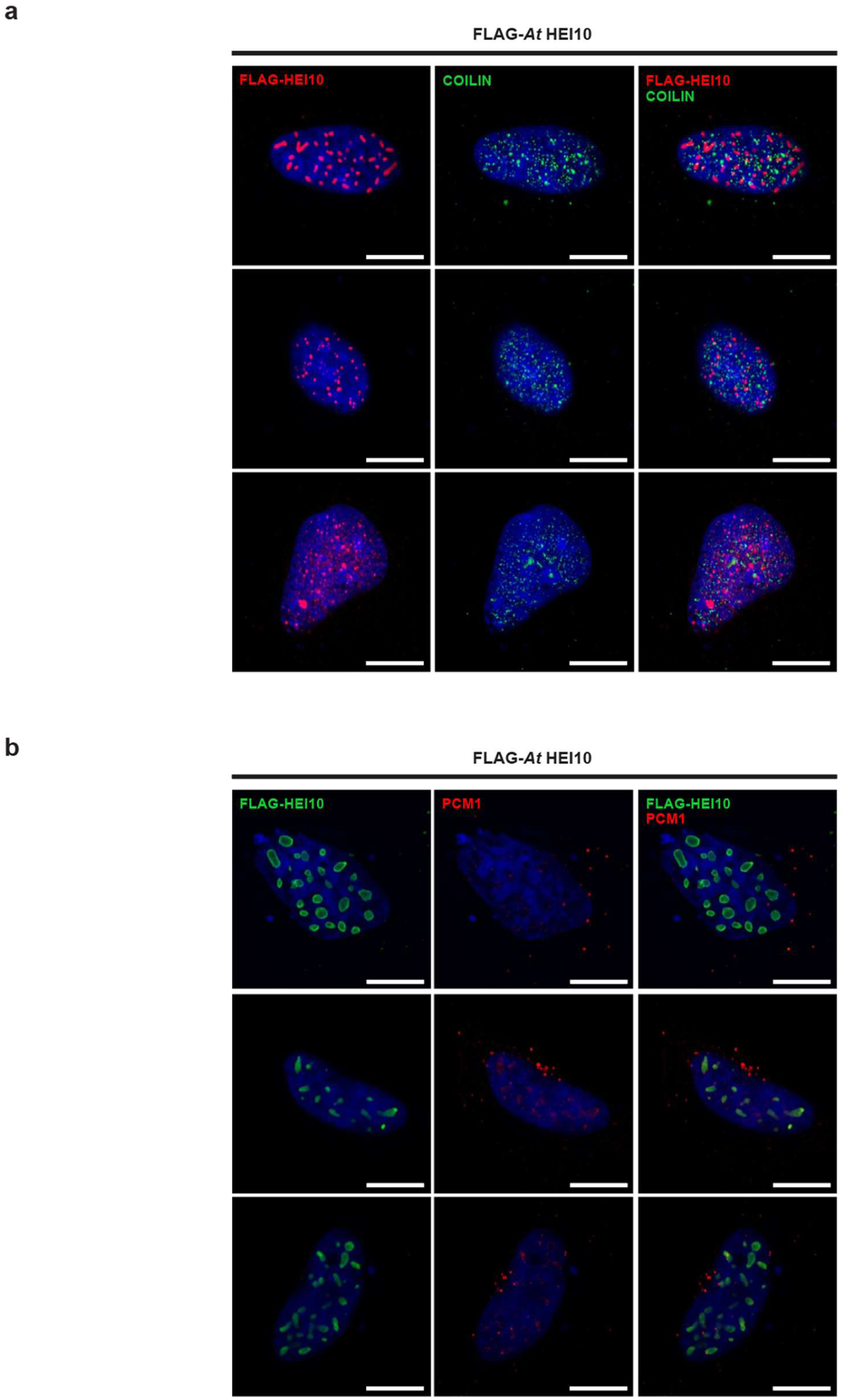
***At* HEI10 foci do not localise with Cajal bodies or peri-centriolar material.** (**a,b**) Flag-*At* HEI10 over-expression in U2OS cells. Co-staining of Flag-*At* HEI10 (red), DAPI (blue), and (**a**) COILIN (green) and (**b**) PCM1 (green). Scale bars, 10 μm.

**Supplementary Figure 15.**
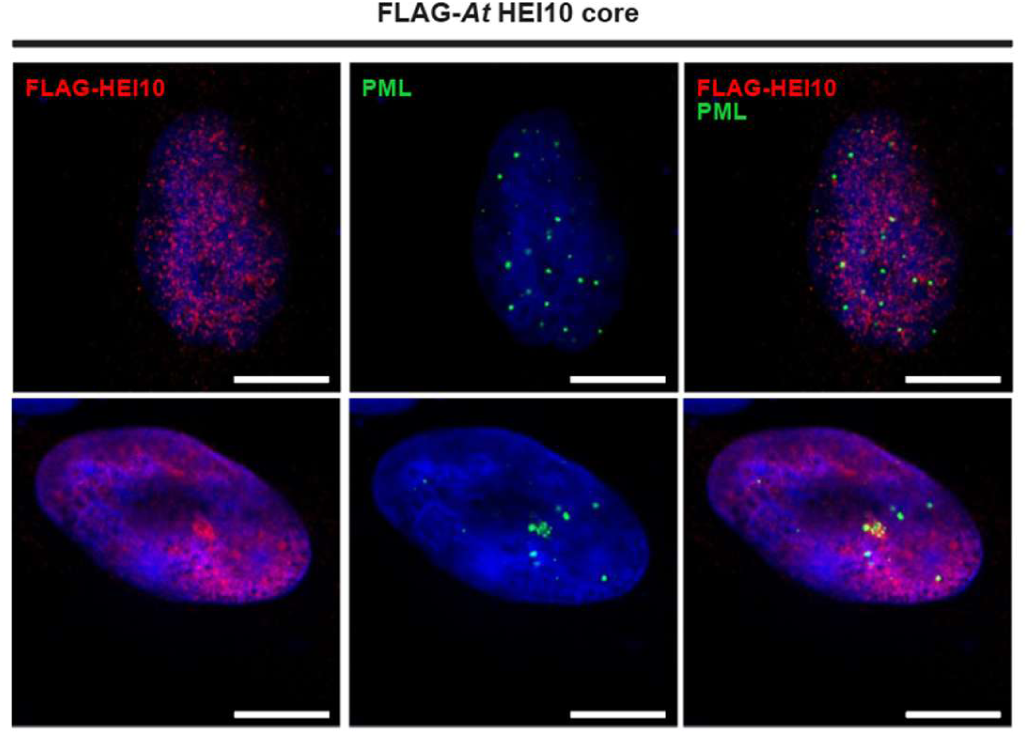
***At* HEI10 core foci do not localise with PML bodies.** Flag-*At* HEI10 over-expression in U2OS cells. Co-staining of Flag- *At* HEI10 (red) PML1 (green) and DAPI (blue), supporting data in Figure 6h. Scale bars, 10 μm.

## Notes

### Competing Interest Statement

The authors have declared no competing interest.

